# Inflammasome activation drives gasdermin-independent plasma membrane rupture by clustering ninjurin-1 in macrophages

**DOI:** 10.64898/2026.04.10.717393

**Authors:** Tadayoshi Karasawa, Hidetoshi Aizawa, Takanori Komada, Yoshiko Mizushina, Emi Aizawa, Chintogtokh Baatarjav, Takahiro Kuchimaru, Yutaka Kodama, Masafumi Takahashi

## Abstract

Inflammasome assembly rapidly triggers caspase-1 activation to initiate pyroptosis, an inflammatory cell death characterized by the release of cytosolic contents, including interleukin (IL)-1β/α. Here, we report that inflammasome activation drives necrotic cell death independent of gasdermin D (GSDMD) and GSDME, which are essential executors of pyroptosis by forming a pore on the plasma membrane and increasing membrane permeability. NLRP3 inflammasome activation induced necrotic cell death, coupled with IL-1β/α release in *Gsdmd*^–/–^*Gsdme*^–/–^ macrophages. Mechanistically, the oligomerization of ninjurin-1 (NINJ1) was caused by inflammasome activation even in the absence of GSDMD and GSDME. Concordantly, glycine, an inhibitor of NINJ1, blocked plasma membrane permeabilization triggered by inflammasome activation in *Gsdmd*^–/–^*Gsdme*^–/–^ macrophages, but not in WT macrophages. The dimerizer-mediated ASC oligomerization promoted NINJ1-mNeonGreen cluster formation in the absence of GSDMD and GSDME. Moreover, NINJ1 deficiency prevented membrane permeabilization initiated by ASC oligomerization in *Gsdmd*^–/–^*Gsdme*^–/–^ immortalized bone marrow-derived macrophages (iBMDM). Blocking of phosphatidylserine (PtdSer) exposure, a feature of inflammasome-driven necrotic cell death, by Xkr8 deficiency inhibited plasma membrane permeabilization in *Gsdmd*^–/–^*Gsdme*^–/–^ iBMDM. These results suggest that inflammasome-triggered activation of caspase-1 itself drives inflammatory necrotic cell death independent of gasdermins.

## Introduction

Upon multiple stresses, cells undergo regulated cell death (RCD), including apoptosis, pyroptosis, necroptosis, and ferroptosis(Newton *et al*, 2024). The mode of cell death is a key determinant of multiple biological processes. In particular, whether cell death occurs via apoptosis or other necrotic cell death is critical for initiating inflammatory responses (Newton *et al*., 2024). While necrotic cell death is accompanied by the release of inflammatory contents as damage/danger-associated molecular patterns (DAMPs), apoptosis preserves plasma membrane integrity and retains cytosolic content in apoptotic bodies. Since apoptotic bodies can be efficiently removed by efferocytosis, apoptosis is regarded as an anti-inflammatory cell death(Doran *et al*, 2020). Therefore, molecular mechanisms that switch the mode of cell death are potential targets for inflammatory diseases.

The inflammasome is a molecular complex that functions as a scaffold for caspase-1 activation(Barnett *et al*, 2023; Karasawa & Takahashi, 2025; Schroder & Tschopp, 2010). Among inflammasomes, nucleotide-binding oligomerization domain, leucine-rich repeat and pyrin domain containing 3 (NLRP3) inflammasome, composed of NLRP3, apoptosis-associated speck-like protein containing a caspase recruitment domain (ASC), and caspase-1, is a key regulator of sterile inflammation because it is activated by various DAMPs, such as extracellular ATP, monosodium urate (MSU) crystals, and cholesterol crystals. Upon inflammasome assembly, the active caspase-1 processes precursors of inflammatory cytokines interleukin (IL)-1β and IL-18 to initiate inflammatory responses. Moreover, the inflammasome complex triggers the activation of multiple caspases, thereby driving diverse cell death cascades. The caspase-1-mediated gasdermin D (GSDMD) processing is a major pathway to initiate pyroptosis, an inflammatory necrotic cell death(Broz *et al*, 2020; Kayagaki *et al*, 2015). Pyroptosis was originally defined as an inflammatory cell death triggered by inflammatory caspases, including caspase-1, 4, and 5(Brennan & Cookson, 2000; Hagar *et al*, 2013). Pyroptosis shares several features with apoptosis, such as DNA fragmentation and phosphatidylserine (PtdSer) exposure. Meanwhile, the rapid induction of increased plasma membrane permeability represents a hallmark of pyroptosis(Brennan & Cookson, 2000). Currently, it is generally accepted that the increased plasma membrane permeability during pyroptosis is caused by the pore formed by GSDMD. Gasdermin family proteins possess pore-forming activities in their amino-terminal domain (NTD)(Ding *et al*, 2016). GSDMD-NTD processed by caspase-1 binds to inositol phospholipid in the plasma membrane and forms an oligomeric pore that results in increased membrane permeability (Liu *et al*, 2016). The pore formed by GSDMD plays critical roles in the release of cytosolic content and inflammatory cytokines IL-1β/α and IL-18 to initiate inflammatory responses(Evavold *et al*, 2018; Monteleone *et al*, 2018). Accumulating evidence suggests that the inflammasome is involved in various inflammatory diseases, including gout, cardiovascular diseases, and neurodegenerative diseases(Barnett *et al*., 2023; Takahashi, 2021). Therefore, inflammasome-triggered GSDMD processing and induction of pyroptosis are expected as a potent therapeutic target for various inflammatory diseases.

Previous studies have suggested that inflammasome assembly leads to activation of caspases other than caspase-1. We have shown that inflammasome activation results in processing of GSDME via activation of caspase-8/3 in the absence of caspase-1(Aizawa *et al*, 2020). GSDME, another gasdermin that is processed by caspase-3, functions as an executor of pyroptosis in a caspase-1-deficient setting. Moreover, multiple studies have suggested that GSDME is processed during inflammasome or caspase-1 activation in the absence of GSDMD(Tsuchiya *et al*, 2019; Zhou & Abbott, 2021). However, it remains unclear whether the blockade of GSDMD and GSDME suffices for the prevention of pyroptosis. To validate the role of GSDMD and GSDME in pyroptosis during inflammasome activation, we established mice lacking both GSDMD and GSDME and found that necrotic cell death occurs even in the absence of GSDMD and GSDME upon inflammasome activation.

## Results

### GSDMD and GSDME are dispensable for necrotic cell death induced by NLRP3 inflammasome activation

Previous studies have suggested that GSDME is involved in the necrotic cell death triggered by inflammasome activation in the absence of GSDMD(Zhou & Abbott, 2021). In order to block gasdermins regulated by the inflammasome, mice lacking both GSDMD and GSDME were developed by crossing *Gsdmd*^–/–^ mice and *Gsdme*^–/–^ mice. As we previously confirmed that necrotic cell death triggered by nigericin in Pam3CSK4-primed primary macrophages was NLRP3 inflammasome dependent(Aizawa *et al*., 2020), we investigated the nigericin-induced cell death in WT, *Casp1/11*^–/–^, *Gsdmd*^–/–^, and *Gsdmd*^–/–^*Gsdme*^–/–^ macrophages. In the early time phase, the combined deficiency of GSDMD and GSDME prevented nigericin-induced pyroptosis (Fig. 1A). However, necrotic cell death occurred in the later time phase of nigericin treatment even in *Gsdmd*^–/–^*Gsdme*^–/–^ macrophages. (Fig. 1 B and C). The onset of inflammasome-triggered necrotic cell death in *Gsdmd*^–/–^*Gsdme*^–/–^ macrophages occurred at an earlier time point than in *Casp1/11*^–/–^ macrophages (Fig. 1C and D). Meanwhile, GSDME deficiency did not affect the kinetics of cell death in *Gsdmd*^–/–^ macrophages. Morphological analysis in live cell imaging showed that cell swelling, a key feature of pyroptosis, occurred in both *Gsdmd*^–/–^ and *Gsdmd*^–/–^*Gsdme*^–/–^ macrophages (Fig. 1D and E).

**Fig. 1.**
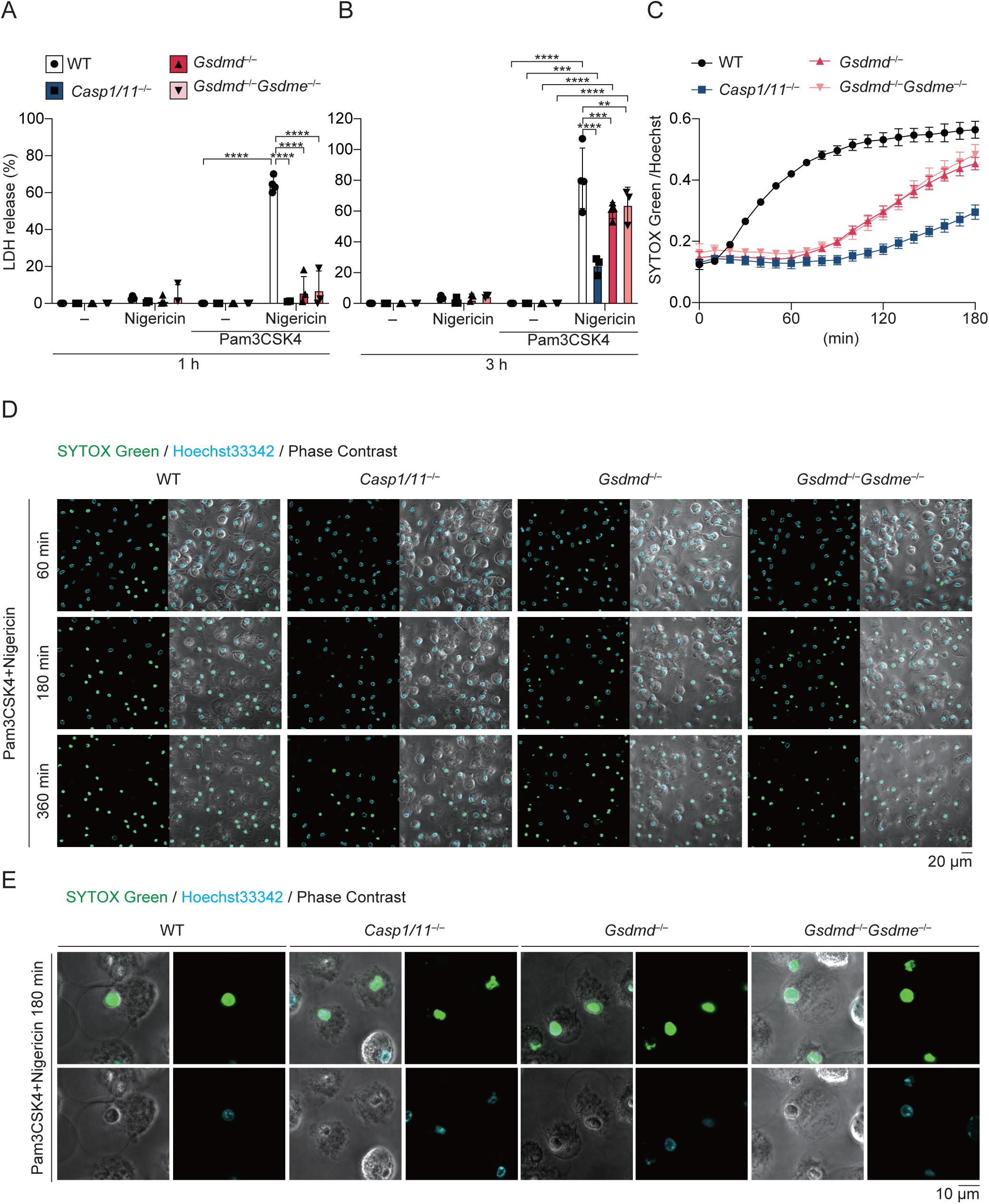
GSDMD and GSDME are dispensable for necrotic cell death induced by NLRP3 activation. (A–E) Primary peritoneal macrophages isolated from WT, *Casp1/11*^–/–^, *Gsdmd*^–/–^, and *Gsdmd*^–/–^*Gsdme*^–/–^ mice were unprimed or primed with Pam3CSK4 (100 ng/mL) for 18 h and then treated with nigericin (5 µM). (A and B) The levels of LDH in the supernatants at 1 h (A) and 3 h (B) were assessed. (C–D) Cells were labelled with Hoechst33342 and then treated with nigericin in the presence of SYTOX Green. (C) Relative fluorescence units of SYTOX Green were measured at 10-min intervals. (D and E) Images were visualized by confocal microscopy. (A and B) The data were obtained from cells derived from 3–4 mice per group. Each dot represents one mouse; the bar indicates mean ± SD. (C) The data were obtained from cells derived from 3 mice per group. Data are shown as mean ± SD. (D and E) Data are representative of at least three independent experiments. (A and B) Statistical significance was calculated using two-way ANOVA with Tukey’s post hoc test. \*\**p* < 0.01, \*\*\**p* < 0.001, \*\*\*\**p* < 0.0001.

### GSDMD and GSDME are dispensable for IL-1 release induced by NLRP3 inflammasome activation

The release of IL-1β is a critical feature of pyroptosis that regulates subsequent inflammatory responses. Recent studies have suggested that the pore formed by GSDMD and GSDME plays a significant role in the release of IL-1β(Evavold *et al*., 2018; Xia *et al*, 2021). Although IL-1β release triggered by NLRP3 inflammasome activation was prevented by GSDMD deficiency in the early time point (Fig. 2A), the IL-1β release was not abrogated in *Gsdmd*^–/–^ macrophages and *Gsdmd*^–/–^*Gsdme*^–/–^ macrophages in the later time point (Fig. 2B). IL-1α, which is dispensable for caspase-1-dependent processing, was also released from *Gsdmd*^–/–^*Gsdme*^–/–^ macrophages in response to inflammasome activation (Fig. 2C and D). Western blot analysis revealed that, during inflammasome activation, the alternative activation of GSDME was potently induced in *Gsdmd*^–/–^ macrophages but was abrogated in *Gsdmd*^–/–^*Gsdme*^–/–^ macrophages (Fig. 2E). The activation of caspase-1 and caspase-3, as well as the processing of IL-1β, were triggered by inflammasome activation in *Gsdmd*^–/–^ and *Gsdmd*^–/–^*Gsdme*^–/–^ macrophages (Fig. 2E and F). Inflammasome-driven necrotic cell death in *Gsdmd*^–/–^*Gsdme*^–/–^ macrophages was mediated by caspase-1 because caspase-1 inhibitor, VX-765, prevented necrotic cell death caused by nigericin-stimulation (Fig. S1A). In agreement with our previous study (Aizawa *et al*., 2020), residual necrotic cell death under caspase-1 inhibition in WT macrophages was prevented by combined deficiency of GSDMD and GSDME. Meanwhile, neither ferroptosis inhibitors (Ferrostatin-1[Fer-1] and liproxistatin-1[Lip-1]) nor necroptosis inhibitor (GSK’872, a RIPK3 inhibitor) inhibited inflammasome-driven necrotic cell death. Moreover, as deficiency of GSDMD and GSDME failed to inhibit inflammasome-mediated IL-1 release in cultured macrophages, we also tested the role of GSDMD and GSDME in inflammatory responses *in vivo*. Indeed, combined deficiency of GSDMD and GSDME did not prevent neutrophil recruitment induced by cholesterol crystals (Fig. 2G and H, Fig. S2), an activator of the inflammasome relevant to atherosclerosis(Duewell *et al*, 2010).

**Fig. 2.**
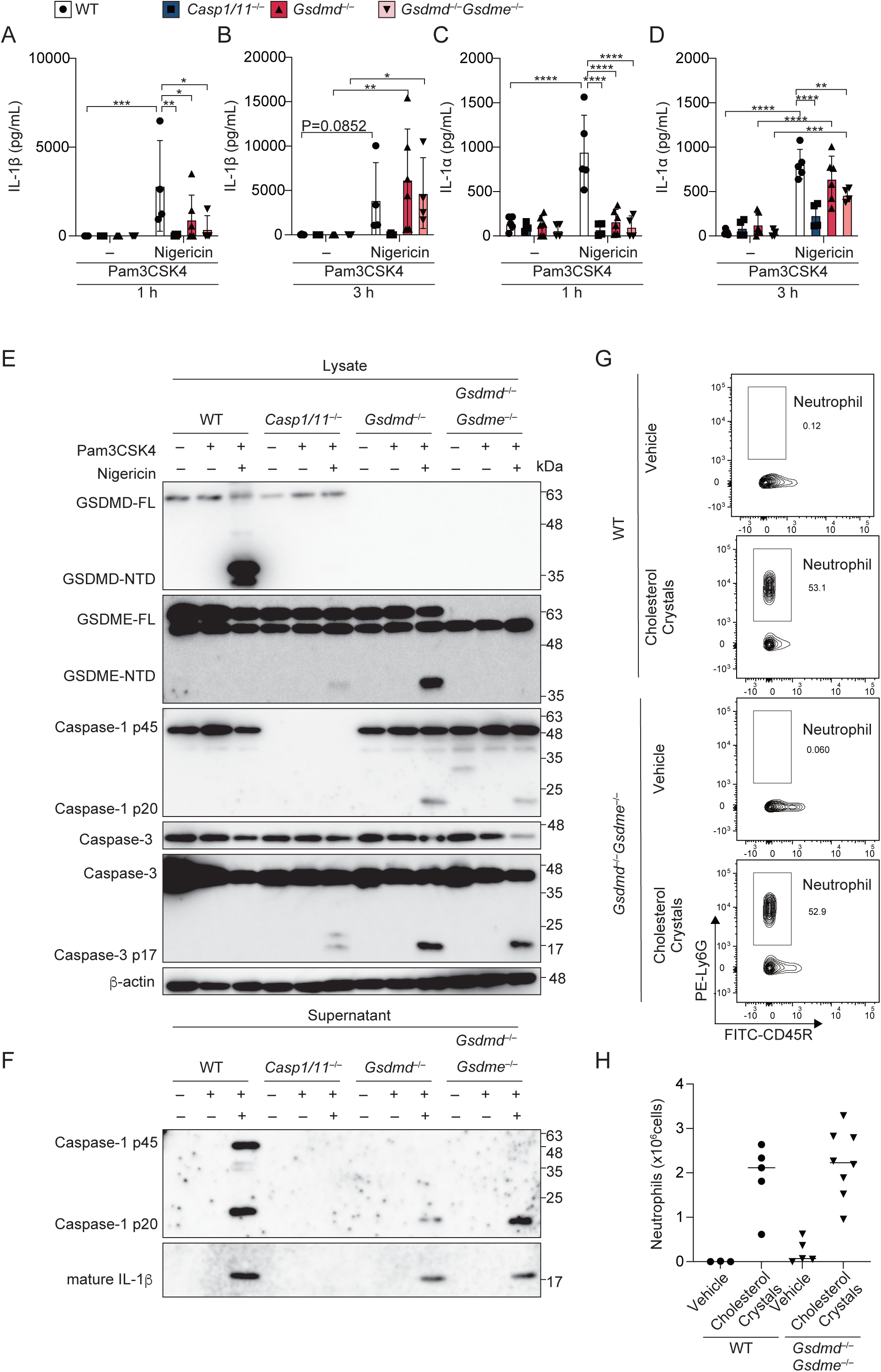
GSDMD and GSDME are dispensable for IL-1 release induced by NLRP3 activation. (A–F) Primary peritoneal macrophages isolated from WT, *Casp1/11*^–/–^, *Gsdmd*^–/–^, and *Gsdmd*^–/–^*Gsdme*^–/–^ mice were unprimed or primed with Pam3CSK4 (100 ng/mL) for 18 h and then treated with nigericin (5 µM) for 1 h (A and C) or 3 h (B, D, E, and F). The levels of IL-1β (A and B) and IL-1α (C and D) in the supernatants were assessed by ELISA. Lysates (E) and supernatants (F) after 3 h stimulation were analyzed by western blot. (G and H) WT and *Gsdmd*^–/–^*Gsdme*^–/–^ mice were intraperitoneally administered cholesterol crystals. After 6 h, peritoneal cells were harvested and analyzed by flow cytometry. (G) Representative plots of neutrophils (CD45^+^Ly6G^+^CD45R^−^). (H) The number of neutrophils in the peritoneal lavage was assessed. (A –D) The data were obtained from cells derived from 4–6 mice per group. Each dot represents one mouse; the bar indicates mean ± SD. (E) Data are representative of at least three independent mice. (G and H) The data were obtained from 3–8 mice per group. (H) Each dot represents one mouse; the bar indicates the median. Statistical significance was calculated using two-way ANOVA with Tukey’s post hoc test. \**p* < 0.05, \*\**p* < 0.01, \*\*\**p* < 0.001, \*\*\*\**p* < 0.0001.

### Gasdermin-independent NINJ1 activation occurs during inflammasome-driven necrotic cell death

To identify the executor of membrane permeabilization during inflammasome-driven necrotic cell death in *Gsdmd*^–/–^*Gsdme*^–/–^ macrophages, we assessed activation of ninjurin-1 (NINJ1), causing membrane rupture (Degen *et al*, 2023; Kayagaki *et al*, 2021). Indeed, the oligomerization of NINJ1 was detected in both nigericin-treated WT and *Gsdmd*^–/–^*Gsdme*^–/–^ macrophages (Fig. 3A). Previous studies have suggested that glycine, a cytoprotectant known to prevent plasma membrane rupture, inhibits NINJ1 activation (Borges *et al*, 2022). In agreement with the previous studies, glycine prevented inflammasome-triggered LDH release but failed to inhibit plasma membrane permeabilization, indicated by SYTOX Green entry in WT macrophages (Fig. 3B and C). By contrast, glycine treatment prevented both LDH release and SYTOX Green entry in *Gsdmd*^–/–^*Gsdme*^–/–^ macrophages (Fig. 3B, D–F). Moreover, the release of IL-1β and IL-1α was blocked by glycine treatment only in *Gsdmd*^–/–^*Gsdme*^–/–^ macrophages (Fig. 3G and H). These results suggest that NINJ1-mediated plasma membrane rupture initiates inflammasome-driven plasma membrane permeabilization and cytokine release in the absence of GSDMD and GSDME.

**Fig. 3.**
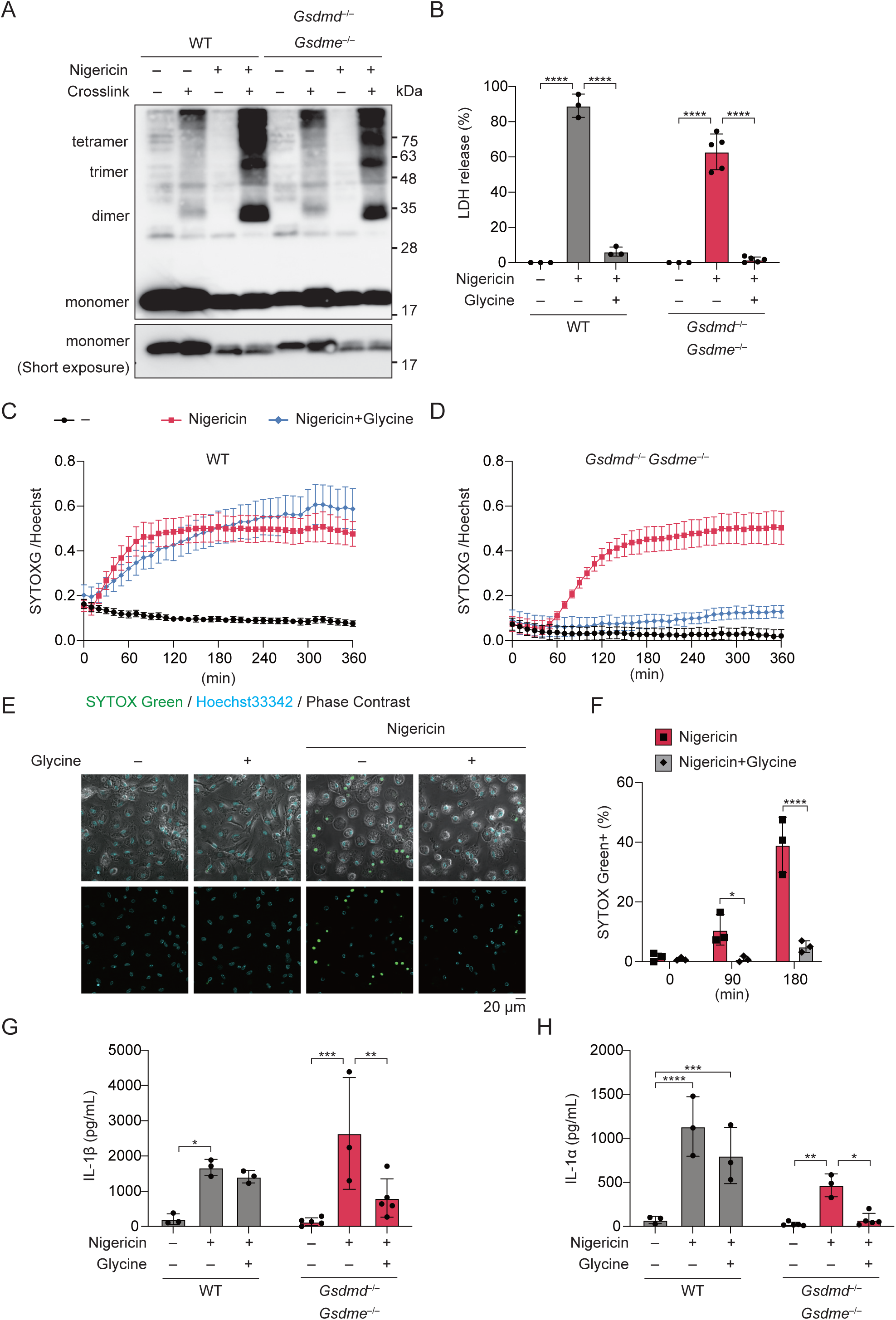
NLRP3 inflammasome activation induces GSDMD- and GSDME-independent NINJ1 assembly. (A) Primary peritoneal macrophages isolated from WT and *Gsdmd*^–/–^*Gsdme*^–/–^ mice were primed with Pam3CSK4 (100 ng/mL) for 18 h and then treated with nigericin (5 µM) for 3 h. Cells were crosslinked with 1.5 % paraformaldehyde and analyzed by western blot. (B –H) Primed WT and *Gsdmd*^–/–^*Gsdme*^–/–^ macrophages were pretreated with 5 mM glycine and then treated with nigericin in the presence (C–F) or absence of SYTOX Green (B, G, and H). (B) The levels of LDH were assessed at 3 h. (C and D) Relative fluorescence units of SYTOX Green were measured at 10-min intervals. (E and F) Images were visualized by confocal microscopy. (E) Representative images. (F) The percentage of SYTOX Green-positive cells was quantified. (G and H) The levels of IL-1β (G) and IL-1α (H) in the supernatants were assessed at 3 h. (B, E–H) The data were obtained from cells derived from 3–5 mice per group. Each dot represents one mouse; the bar indicates mean ± SD. (C and D) The data were obtained from cells derived from 3 mice per group. Data are shown as mean ± SD. Statistical significance was calculated using two-way ANOVA with Tukey’s post hoc test. \**p* < 0.05, \*\**p* < 0.01, \*\*\**p* < 0.001, \*\*\*\**p* < 0.0001.

### Inhibition of NINJ1 prevents inflammasome-driven necrotic cell death in the absence of GSDMD and GSDME

To mimic inflammasome-driven necrotic cell death, immortalized bone marrow-derived macrophages (iBMDM) were transduced with the fusion protein consisting of mutated FKBP12 and ASC (*DmrBASC-*iBMDM, Fig. S3A). Then, *DmrBASC-*iBMDM were transduced with tandem gRNA targeting *Gsdmd* and *Gsdme*, and enCas12a to develop *Gsdmd/e* double-knockout (DKO) iBMDM (Fig. S3B and S3C). ASC oligomerization-triggered necrotic cell death in *DmrBASC-Gsdmd/e* DKO iBMDM was caspase-1-dependent (Fig. S3D). Moreover, rapid loss of cytosolic content during necrotic cell death was visualized by transduction of hKusabira Orange 1 fluorescent protein (Fig. S3E and F). Consistent with the observation in primary macrophages, LDH release and increased plasma membrane permeability induced by ASC oligomerization were prevented by the treatment with glycine in *DmrBASC-Gsdmd/e* DKO iBMDM (Fig. 4A–C). By contrast, in the presence of GSDMD and GSDME, whereas LDH release induced by ASC oligomerization was prevented by the treatment with glycine, increased plasma membrane permeability was not prevented by glycine treatment (Fig. S3G and H). To assess the involvement of NINJ1 in increased plasma membrane permeability, *DmrBASC-Gsdmd/e* DKO iBMDM were transduced with a lentiviral vector expressing NINJ1-mNeonGreen fusion protein under TRE promoter. NINJ1-mNeonGreen was localized throughout the plasma membrane in steady states. Upon induction of ASC oligomerization, NINJ1-mNeonGreen forms clusters on the plasma membrane in necrotic cells (Fig. 4D). To further assess the involvement of NINJ1 in inflammasome-driven necrotic cell death, gRNA targeting *Ninj1* and Cas9 were transduced to *DmrBASC-Gsdmd/e* DKO iBMDM (Fig. S3J). Consistently, NINJ1-deficiency prevented ASC oligomerization-triggered plasma membrane rupture and permeabilization (Fig. 4E–G and Fig. S3K).

**Fig. 4.**
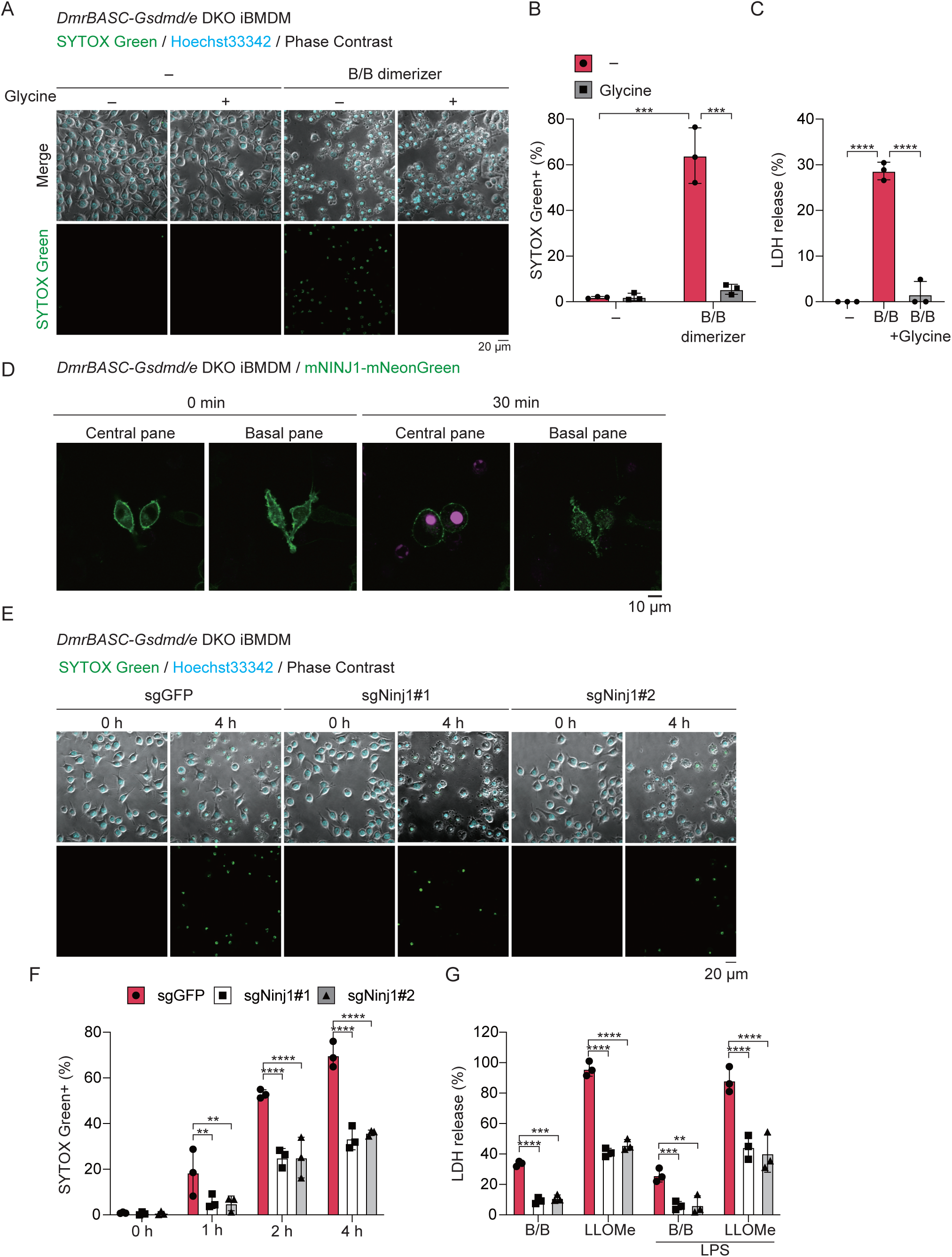
NINJ1-deficiency prevents necrotic cell death triggered by ASC oligomerization in the absence of GSDMD and GSDME. (A–C) *DmrBASC-Gsdmd/e* DKO iBMDM were pretreated with 5 mM glycine and then treated with 100 nM B/B dimerizer in the presence (A and B) or absence of SYTOX Green (C). (A and B) Images were visualized by confocal microscopy. (A) Representative images. (B) The percentage of SYTOX Green-positive cells was quantified. (C) The levels of LDH were assessed at 4 h. (D) *DmrBASC-TRE-NINJ1mNeonGreen–Gsdmd/e* DKO iBMDM were treated with 100 nM B/B dimerizer in the presence of SYTOX Deep red. Images were visualized by confocal microscopy. (E–G) *DmrBASC-Gsdmd/e* DKO iBMDM were transduced with LentiCRISPRv2 expressing GFP or Ninj1-targeted gRNA and treated with 100 nM B/B dimerizer. (E and F) Cells were stimulated in the presence of SYTOX Green, (E) Images were visualized by confocal microscopy. (F) The ratio of SYTOX Green-positive cells was quantified. (G) Cells were treated with 100 nM B/B dimerizer or LLOMe. The levels of LDH were assessed at 4 h. (A–F) The data were obtained from 3 independent experiments. (B, C, F, and G) Each dot represents independent experiments; bar indicates mean ± SD. (A, D, and E) The data are representative of 3 independent experiments. Statistical significance was calculated using two-way ANOVA with Tukey’s post hoc test. \*\**p* < 0.01, \*\*\**p* < 0.001, \*\*\*\**p* < 0.0001.

### Exposure of phosphatidyl serine precedes or coincides with plasma membrane permeabilization

To explore the mechanisms of NINJ1-mediated cell death rupture, we focused on the changes in the plasma membrane during cell death. Exposure of phosphatidyl serine (PtdSer) is a well-known feature of caspase-initiated apoptosis and pyroptosis(Miao *et al*, 2011). We speculated that the exposure of PtdSer is involved in NINJ1 activation because exposure of PtdSer was increased in B/B-dimerizer-treated NINJ1-deficient *DmrBASC-Gsdmd/e* DKO iBMDM (Fig. S3K). Live cell imaging of PtdSer exposure and membrane permeability was assessed using FITC-annexin V and SYTOX Deep red in nigericin-stimulated *Gsdmd*^–/–^*Gsdme*^–/–^ macrophages. Exposure of PtdSer and entry of SYTOX Deep red occurred in two modes; PtdSer exposure preceded or coincided with plasma membrane permeabilization (Fig. 5A–F, Fig. S4A and B). Furthermore, the increased Annexin V+ 7AAD+ cells during ASC oligomerization were confirmed in *DmrBASC-Gsdmd/e* DKO iBMDM (Fig. 5G–I). Of note, Annexin V–7-AAD+ cells were not detected. On the other hand, cell swelling is a key feature of pyroptosis, and necrotic *Gsdmd*^–/–^*Gsdme*^–/–^ macrophages exhibit swollen morphologies (Fig. 1E). Therefore, we evaluated the volume of the cell before membrane rupture. Although an increasing trend of cell sphericity was detected, cell volume was unchanged during the induction of inflammasome-driven necrotic cell death in *Gsdmd*^–/–^*Gsdme*^–/–^ macrophages (Fig. 5J–O). Similar results were observed in *DmrBASC-Gsdmd/e* DKO iBMDM (Fig. S4 C–I). Moreover, plasma membrane tension assessed by Flipper-TR was not increased during inflammasome activation (Fig. S4J and K). These results indicate that increased cell volume and membrane tension are not the cause of NINJ1 activation in inflammasome-driven necrotic cell death in *Gsdmd*^–/–^*Gsdme*^–/–^ macrophages.

**Fig. 5.**
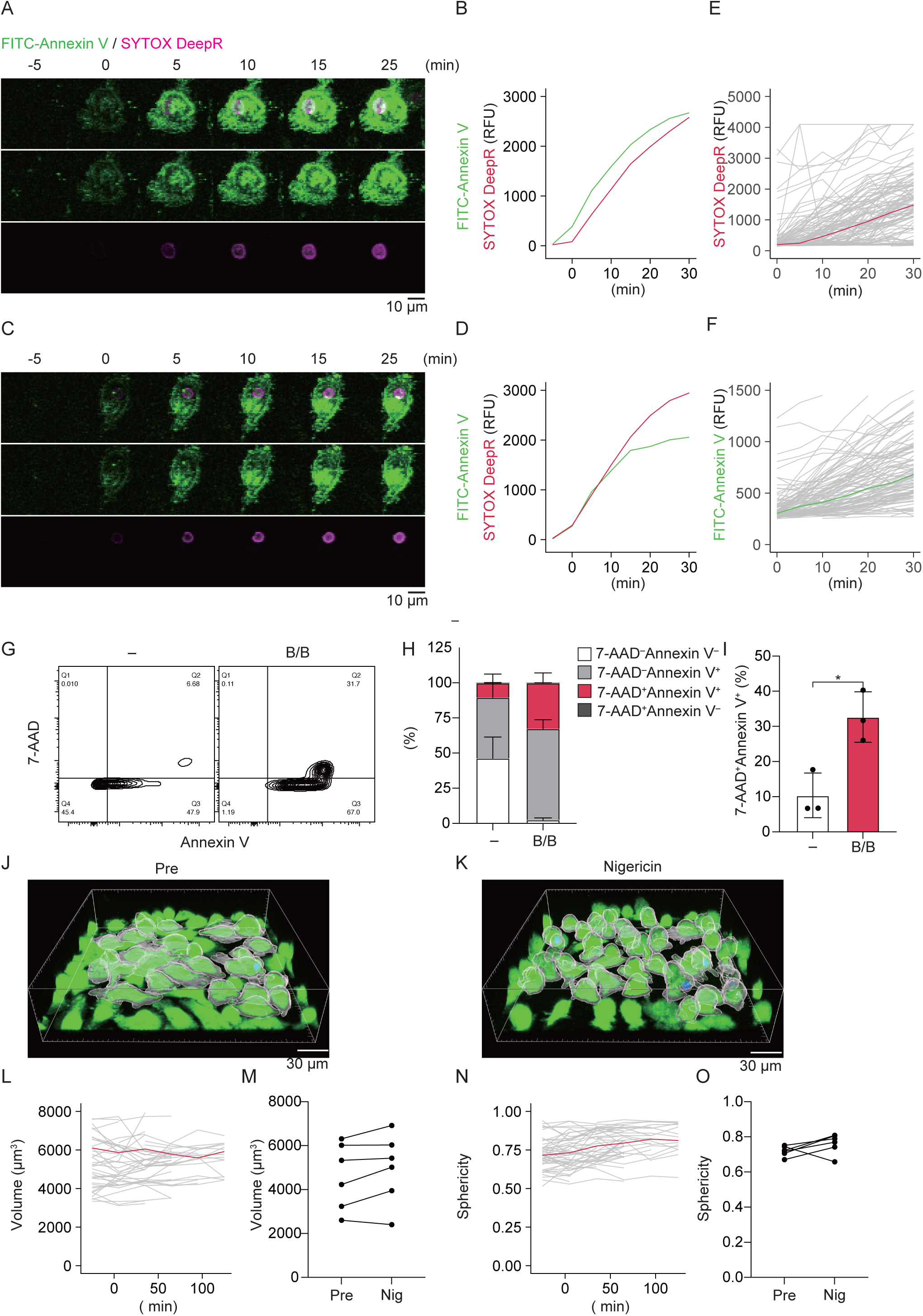
Exposure of phosphatidyl serine during inflammasome-driven necrotic cell death. (A–F) Primary peritoneal macrophages isolated from *Gsdmd*^–/–^*Gsdme*^–/–^ mice were primed with Pam3CSK4 (100 ng/mL) for overnight and treated with 5 µM nigericin in the presence of FITC-Annexin V and SYTOX Deep red. Z-stack time-lapse imaging was performed at 5 min-intervals. Annexin V-positive cells were tracked and analyzed. T_0_ is defined as a time frame in which FITC-Annexin V signals were detected. (A and B) Representative cell in which exposure of PtdSer precedes the plasma membrane permeabilization. (A) Montage images and (B) quantification of fluorescent signals. (C and D) Representative cell in which exposure of PtdSer coincides with plasma membrane permeabilization. (C) Montage images and (D) quantification of fluorescent signals. (E and F) Representative plots of SYTOX Deep red (E) and FITC-Annexin V (F). Gray lines indicate individual cells, and red line (E) or green line (F) indicates the median of all cells obtained from a single mouse. (G–I) *DmrBASC-Gsdmd/e* DKO iBMDM were treated with100 nM B/B dimerizer for 4 h. Cells were stained with FITC-Annexin V and 7-AAD and analyzed by flow cytometry. (G) Representative plot, (H) the percentage of each population, and (I) the percentage of Annexin V^+^7-AAD^+^ cells. Each dot represents independent experiments; the bar indicates mean ± SD. (J–O) Primed *Gsdmd*^–/–^*Gsdme*^–/–^ macrophages were stained with 5 µM CFSE and then treated with 5 µM nigericin in the presence of SYTOX Deep red. Z-stack time-lapse imaging was performed at 30-min intervals. (J and K) Representative 3D images of cells (J) before stimulation and 60 min after nigericin treatment (K). (L–O) The cell volume and sphericity in live cells were tracked. Representative plots of volume (L) and sphericity (N). Gray lines indicate individual cells, and the red line indicates the mean of all cells obtained from a single mouse. (M and O) Each dot represents the mean values of cells derived from individual mice. Cell volume and sphericity were assessed at one frame before the first detection of SYTOX Deep red-positive cells. (A–F, J–O) The data were obtained from cells derived from 6 mice. (G–I) The data are obtained from 3 independent experiments. Statistical significance was calculated using Student’s t test. \**p* < 0.05

### Xkr8-dependent PtdSer exposure is essential for inflammasome-driven necrotic cell death in the absence of GSDMD and GSDME

Multiple phospholipid scramblases are known to translocate PtdSer. Xkr8 is known as a caspase-dependent scramblase that is involved in PtdSer exposure during apoptosis (Suzuki *et al*, 2013). On the other hand, Ano6 (also known as TMEM16F) functions as a Ca^2+^-dependent phospholipid scramblase (Suzuki *et al*, 2010). Tmem63b is also reported as a membrane structure-responsive lipid scramblase (Miyata *et al*, 2025). To further investigate the involvement of PtdSer exposure in inflammasome-driven necrotic cell death in the absence of GSDMD and GSDME, gRNA targeting phospholipid scramblases were transduced into *DmrBASC-Gsdmd/e* DKO iBMDM. The mutations were successfully introduced by all employed gRNAs (Fig. S5A–F). Among these phospholipid scramblases, gRNA-targeting *Xkr8* attenuated SYTOX entry upon ASC oligomerization (Fig. 6A and B). The release of LDH was also attenuated by the transduction of *Xkr8*-targeting gRNA (Fig. 6C). Although 2 gRNAs were designed for *Xkr8*, *Xkr8*#1 showed lower frequency of frameshift mutation than *Xkr8*#2 (Fig. S5A and B). Meanwhile, *Xkr8*#2 more efficiently prevented ASC oligomerization-triggered PtdSer exposure and prevented induction of necrotic cell death (Fig. 6D–F and Fig. S6B–D). On the other hand, transduction of gRNA targeting *Ano6* and *Tmem63b* failed to inhibit PtdSer exposure and necrotic cell death (Fig. 6A–C, E, F, and Fig. S6A–D)

**Fig. 6.**
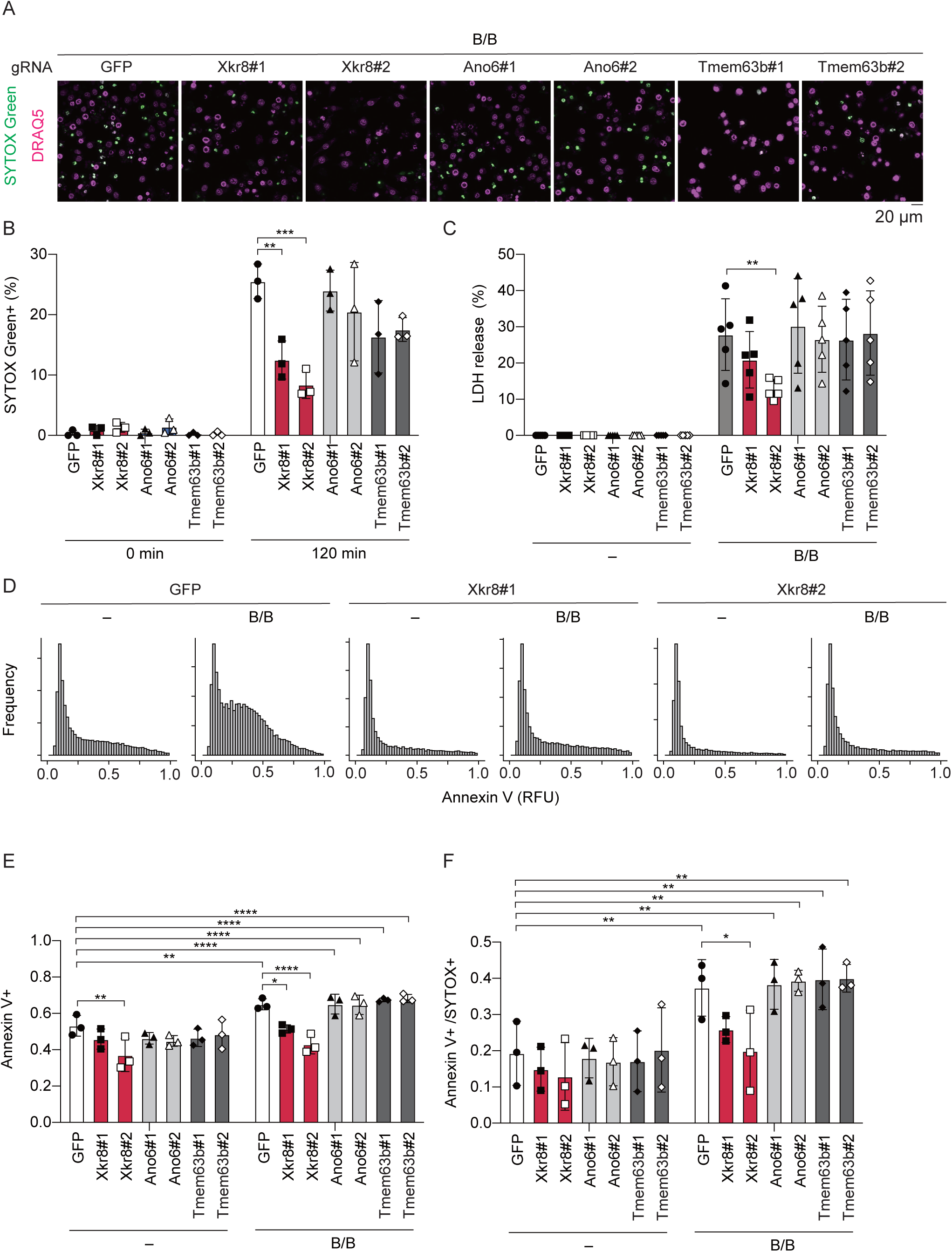
Xkr8-dependent PtdSer exposure is essential for necrotic cell death triggered by ASC oligomerization in the absence of GSDMD and GSDME. (A–F) *DmrBASC-Gsdmd/e* DKO iBMDM were transduced with LentiCRISPRv2 expressing GFP, Xkr8, Ano6, and Tmem63b-targeted gRNA. The cells were treated with 100 nM B/B dimerizer. (A and B) Cells were stimulated in the presence of SYTOX Green. (A) Images were visualized by confocal microscopy at 2h. (B) The percentage of SYTOX Green-positive cells was quantified. (C) The levels of LDH were assessed at 4 h. (D–F) Cells were stimulated in the presence of FITC-Annexin V and SYTOX Deep red for 2 h and analyzed by high content analysis. (D) Representative histogram of Annexin V intensity. (E) The ratio of Annexin V+ cells and Annexin V+ SYTOX Deep red+ cells was quantified. (A–F) The data were obtained from 3 (A, B, D–F) or 5 (C) independent experiments. Each dot represents individual experiments; the bar indicates mean ± SD. Statistical significance was calculated using two-way ANOVA with Tukey’s post hoc test. \**p* < 0.05, \*\**p* < 0.01, \*\*\**p* < 0.001, \*\*\*\**p* < 0.0001.

## Discussion

Inflammasome activation engages multiple pathways that trigger necrotic cell death to initiate inflammatory responses. In the present study, we found that combined deficiency of GSDMD and GSDME failed to prevent necrotic cell death triggered by inflammasome activation. This necrotic cell death is accompanied by the release of pyroptosis-associated cytokines, IL-1β and IL-1α. Concordantly, deficiency of GSDMD and GSDME failed to inhibit infiltration of neutrophils induced by cholesterol crystals in mice. Furthermore, the inflammasome promotes NINJ1 oligomerization in the absence of GSDMD and GSDME. Deficiency of NINJ1 prevents not only membrane rupture but also membrane permeabilization upon inflammasome activation. Among the features associated with inflammasome-driven cell death, exposure of PtdSer precedes or coincides with plasma membrane permeabilization. The exposure of PtdSer plays a causative role because the deficiency of Xkr8 attenuated the exposure of PtdSer and membrane permeabilization during necrotic cell death triggered by inflammasome activation in the absence of GSDMD and GSDME. Our data clearly suggest that GSDMD and GSDME are dispensable for the induction of NINJ1-mediated plasma membrane rupture triggered by inflammasome activation.

Previous studies have suggested that a deficiency of GSDMD is insufficient for the prevention of inflammasome-dependent necrotic cell death(Kayagaki *et al*., 2015; Schneider *et al*, 2017). While caspase-11-initiated pyroptosis is completely blocked by the deficiency of GSDMD, canonical inflammasome-triggered necrotic cell death has not been attributed to a single executor. Zhou *et al*. reported that GSDME is alternatively processed during NLRP3 inflammasome activation in the GSDMD-deficient THP-1 cells(Zhou & Abbott, 2021). The processed GSDME mediates IL-1β release and secondary pyroptosis. However, Broz and his colleagues(Heilig *et al*, 2020) reported that GSDME has no contribution to AIM2 inflammasome-driven necrotic cell death in GSDMD-deficient iBMDM. As previous studies have investigated the involvement of GSDMD and GSDME in cell lines, we developed mice lacking GSDMD and GSDME and confirmed that the deficiency of GSDMD and GSDME is dispensable for the induction of necrotic cell death triggered by the inflammasome in primary macrophages.

Pyroptosis was originally described as a caspase-1-dependent necrotic cell death in *Salmonella*-infected macrophages(Brennan & Cookson, 2000), and had been regarded as a cell death mediated by inflammatory caspases. Since the identification of GSDMD as a pore-forming protein processed by inflammatory caspases(Kayagaki *et al*., 2015; Shi *et al*, 2015), pyroptosis has been defined as a gasdermin-dependent cell death (Broz *et al*., 2020). In the present study, in the absence of GSDMD and GSDME, inflammasome activation drives necrotic cell death that shares similar features with pyroptosis. As *Gsdmd*^–/–^*Gsdme*^–/–^ macrophages released a comparable amount of IL-1β and IL-1α upon NLRP3 inflammasome activation, we consider that GSDMD determines the kinetics of cell death induction rather than the secretome of inflammasome-triggered necrotic cell death. Notably, the necrotic cell death found in this study is dependent on caspase-1. Thus, we postulate that caspase-1 is a critical determinant of the pyroptotic feature in inflammasome-triggered necrotic cell death.

NINJ1 was reported as an executor of plasma membrane rupture in the final phase of pyroptosis(Kayagaki *et al*., 2021). Subsequent studies have suggested that NINJ1 is involved in plasma membrane rupture in multiple modes of RCD(Dondelinger *et al*, 2023; Ramos *et al*, 2024). In the present study, we found that membrane permeabilization as well as plasma membrane rupture was prevented by NINJ1-deficiency in inflammasome-triggered necrotic cell death in the absence of GSDMD and GSDME. Regarding NINJ1-mediated plasma membrane permeabilization, Ramos *et al*. reported that NINJ1 regulates both membrane permeabilization and plasma membrane rupture in ferroptosis(Ramos *et al*., 2024). Since ferroptosis is accompanied by the peroxidation of phospholipid in the plasma membrane, it is plausible that NINJ1 triggers plasma membrane permeabilization in response to the changes in the condition of the plasma membrane. With regard to the condition of plasma membrane and NINJ1 activation, a recent study demonstrated that NINJ1 is involved in mechanical strain-induced plasma membrane rupture(Zhu *et al*, 2025). However, we did not detect an increase in plasma membrane tension in inflammasome-triggered necrotic cell death. Although cell swelling has also been reported as a key determinant of NINJ1 activation(Dondelinger *et al*., 2023), the volume of cells was not altered in our study.

Necrotic cell death observed in the present study is dependent on caspase-1. Therefore, we focused on the PtdSer exposure, a well-established feature of caspase-dependent RCD. An asymmetric lipid distribution within the lipid bilayer is regulated by various enzymes. Flippases and floppases translocate specific lipids from the outer to the inner leaflet or from the inner to the outer leaflet(Nagata, 2018). Thus, PtdSer is tightly kept in the inner leaflet of the plasma membrane by ATP-dependent flippases in steady states. In the present study, we observed that PtdSer exposure precedes or coincides with plasma membrane permeabilization triggered by the inflammasome in the absence of GSDMD and GSDME. Among several scramblases involved in PtdSer translocation, Xkr8, a caspase-dependent phospholipid scramblase, was involved in PtdSer exposure and membrane permeabilization in *Gsdmd*^–/–^*Gsdme*^–/–^ macrophages. Previous studies reported that Ca^2+^-dependent phospholipid scramblase, Ano6 mediates PtdSer exposure in response to GSDMD-pore-dependent Ca^2+^ mobilization(Ousingsawat *et al*, 2018; Yang *et al*, 2019). However, Ano6 deficiency did not affect PtdSer exposure in the absence of GSDMD and GSDME. Hence, deficiency of Xkr8 seems to be sufficient to block PtdSer exposure in the GSDMD- and GSDME-deficient settings. Although further investigations are needed to clarify the link between NINJ1 assembly and Xkr8-dependent PtdSer exposure, a previous study suggested that NINJ1 is associated with phospholipid in the outer leaflet of the plasma membrane(Sahoo *et al*, 2025). Accordingly, it is possible that the altered lipid distribution in the plasma membrane regulates NINJ1 assembly upon caspase activation. Meanwhile, Xkr8 is activated by cleavage of caspase-3 or caspase-7 and is involved in the exposure of PtdSer during apoptosis. Although we assessed only inflammasome-triggered plasma membrane rupture, it is plausible that Xkr8-mediated PtdSer exposure mediates broad plasma membrane rupture during secondary necrosis.

In the present study, although we investigated the mechanisms of necrotic cell death triggered by inflammasome activation in the absence of GSDMD and GSDME, there are several limitations. First, we investigated the effect of NINJ1-deficiency only in cultured iBMDM, whose characteristics are distinct from primary macrophages; therefore, further investigation into the contribution of NINJ1 using NINJ1-deficient mice is required. Second, although we developed iBMDM expressing NINJ1-mNeonGreen reporter to analyze NINJ1 activation during necrotic cell death, we could not investigate the detailed time course of NINJ1 activation due to the cell vulnerability to phototoxicity. Therefore, seamless time-lapse imaging with reduced phototoxicity is required to evaluate Xkr8-mediated PtdSer exposure on NINJ1 assembly. Further assessment using improved experimental settings will provide valuable information on the mechanisms of NINJ1 assembly.

In conclusion, we found that inflammasome activation can initiate necrotic cell death via the gasdermin-independent but NINJ1-dependent mechanism. Our findings will be valuable for the development of novel therapeutic strategies for inflammasome-associated inflammatory diseases.

## Materials and Methods

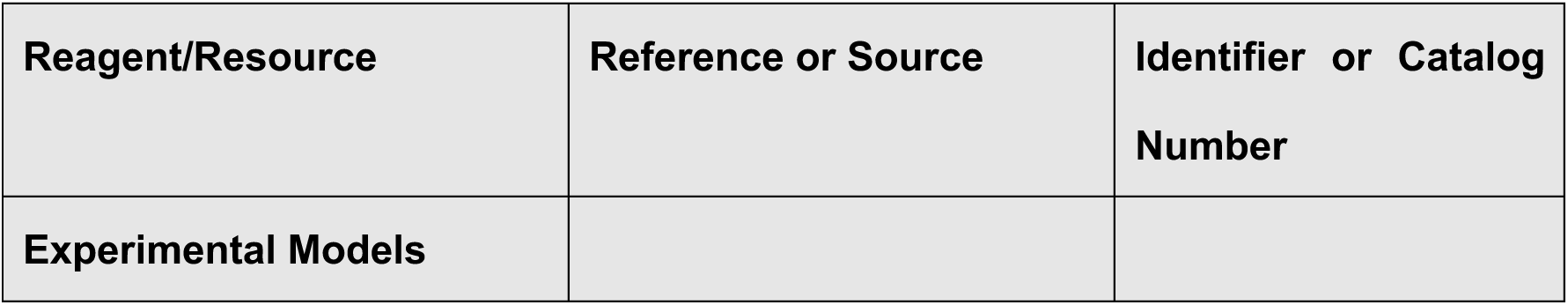
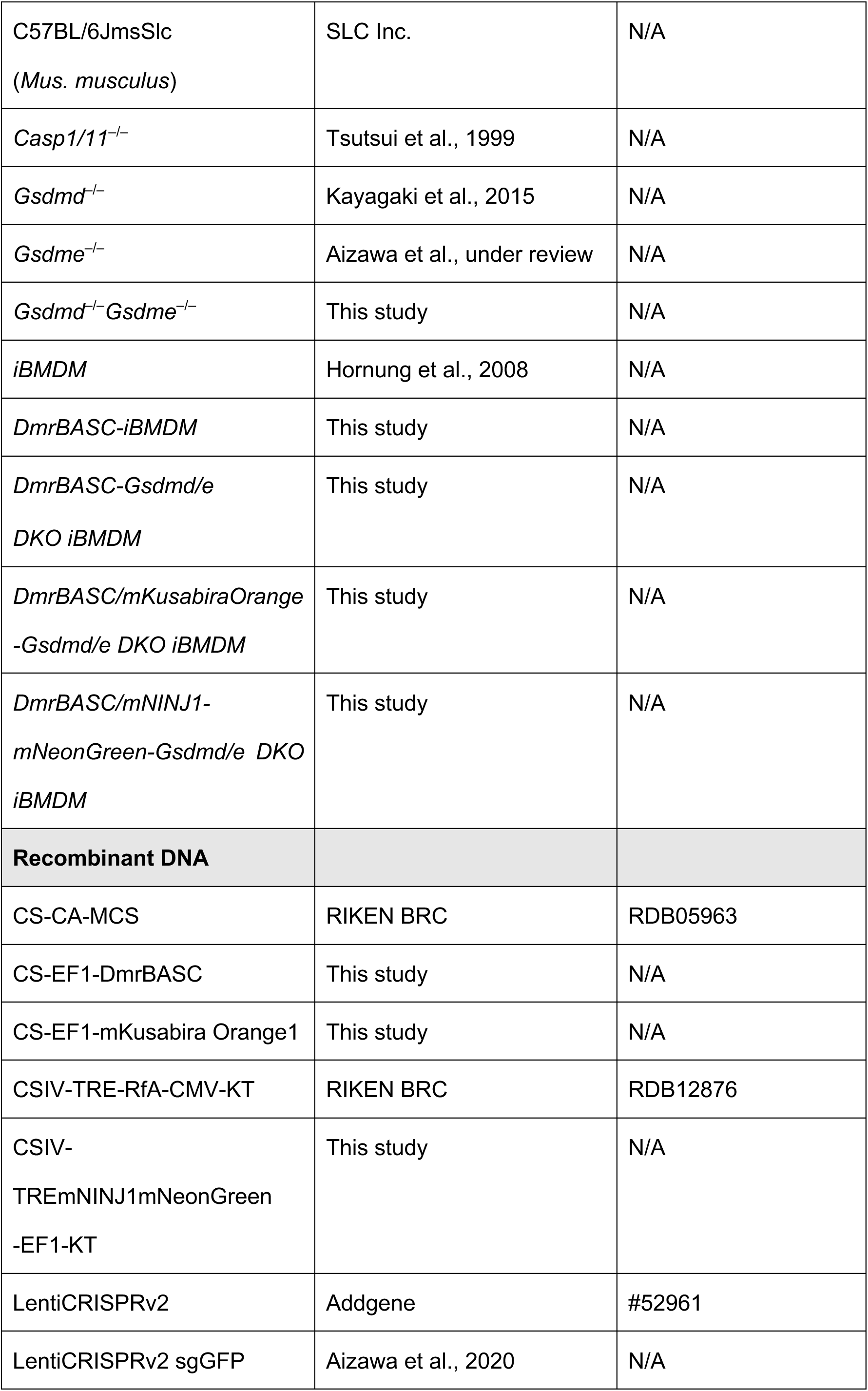
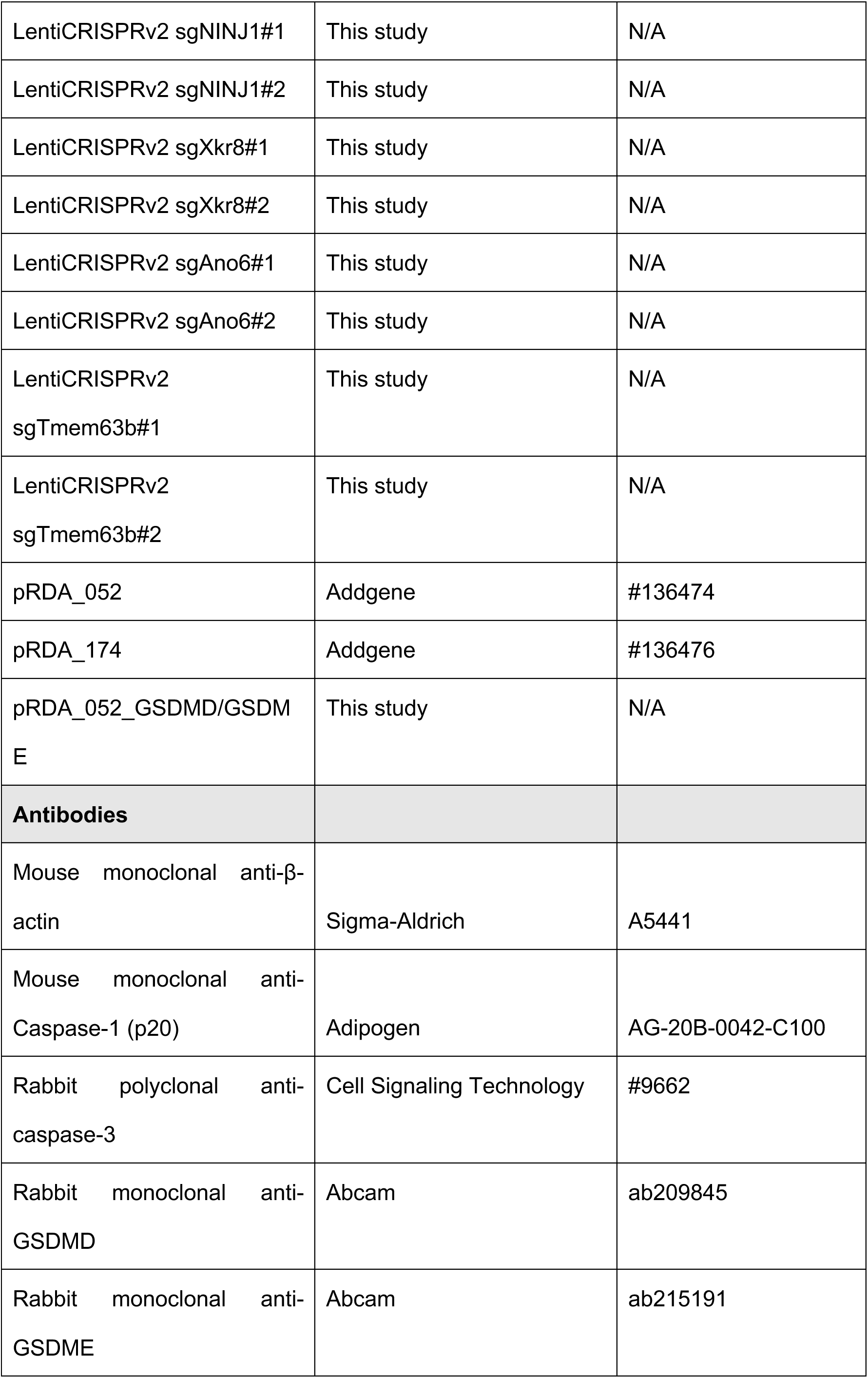
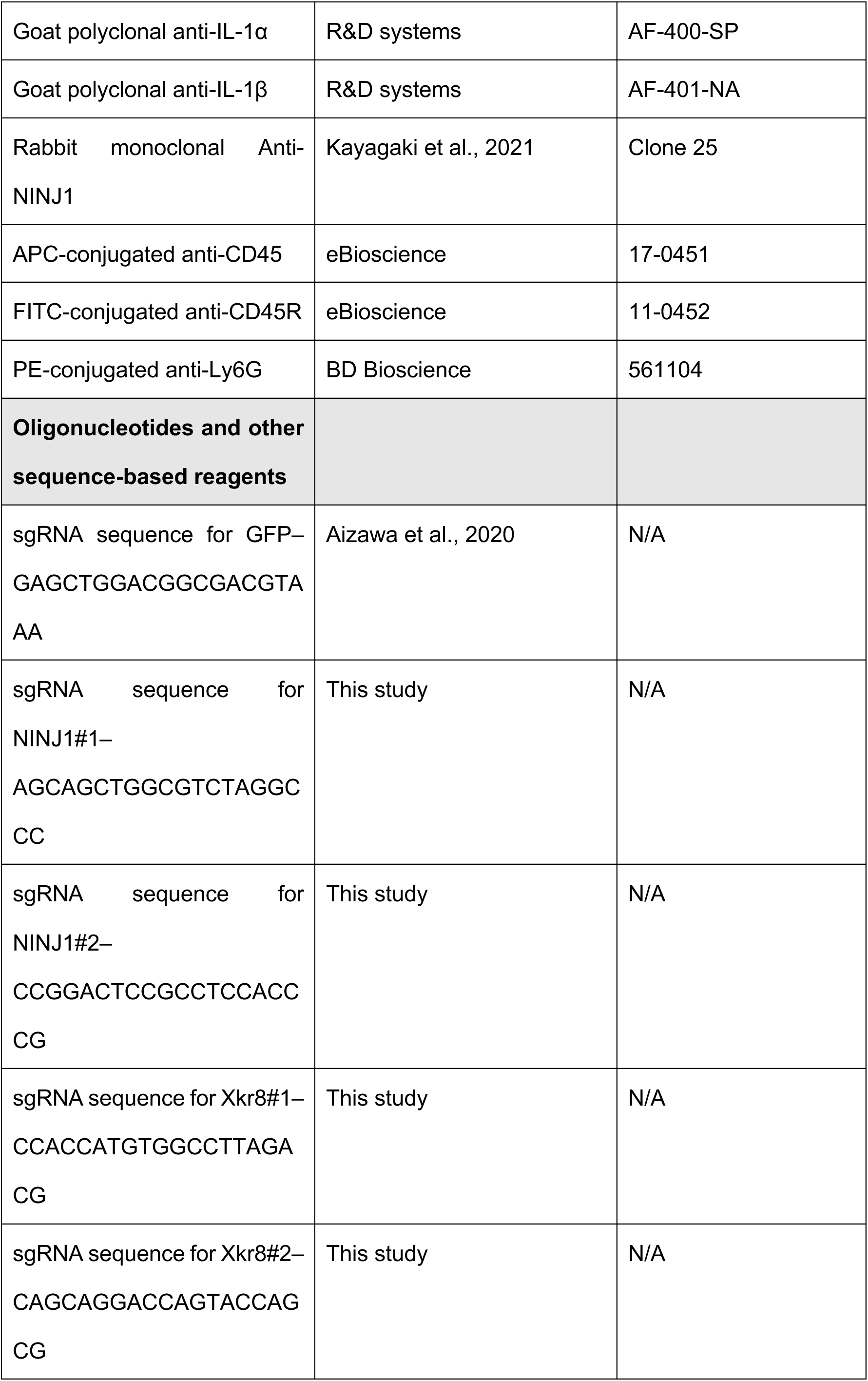
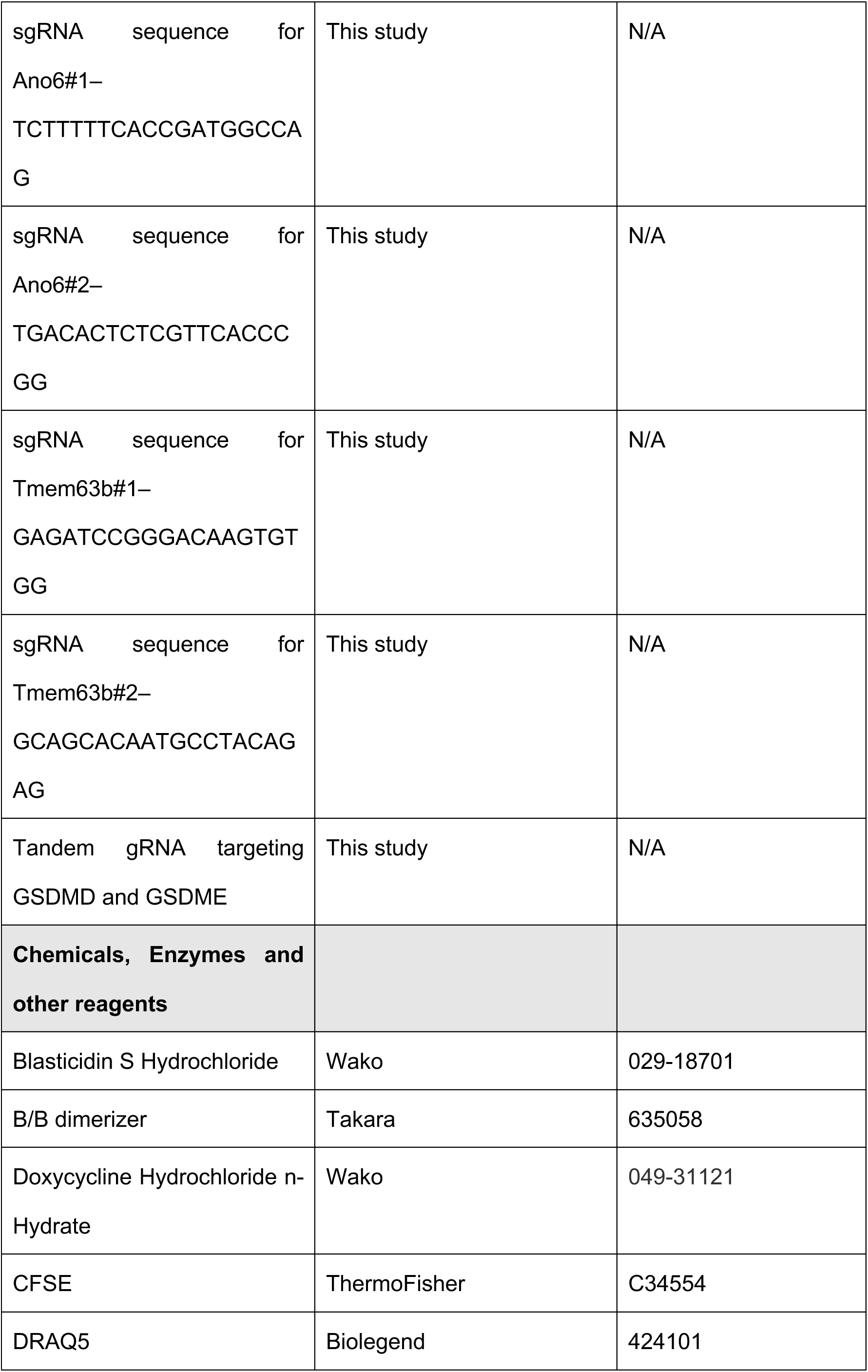
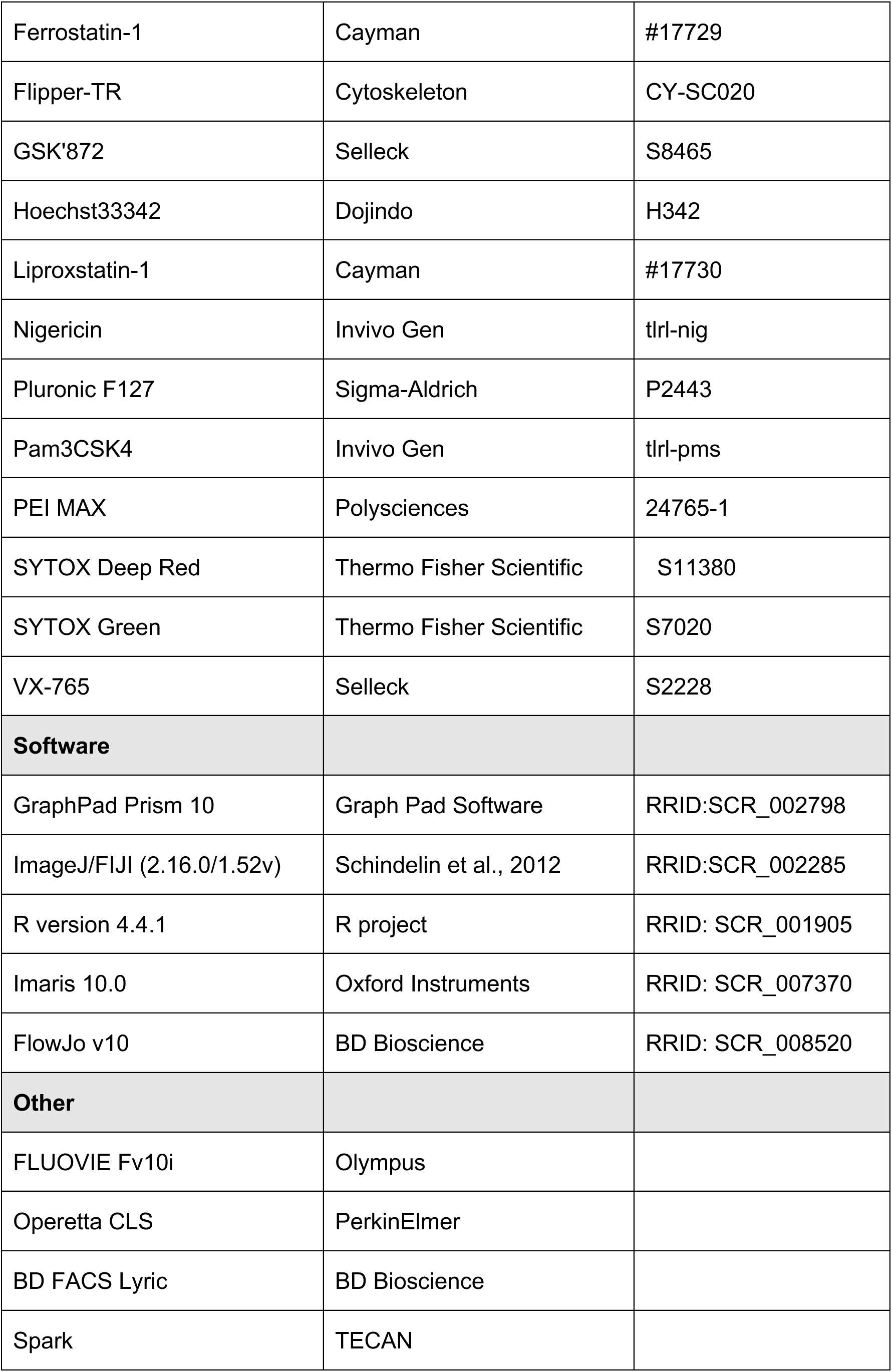

### Animals and primary culture

C57BL/6J mice were purchased from SLC Inc. (Shizuoka, Japan). *Casp1/11*^–/–^ were provided by Dr. Hiroko Tsusui Hyogo College of Medicine, Nishinomiya, Japan). (Tsutsui *et al*, 1999). *Gsdmd*^–/–^ mice were developed by Genentech (South San Francisco, CA), (Kayagaki *et al*., 2015) and provided by Dr. Kate Schroder (University of Queensland, Australia). *Gsdme*^–/–^ mice were generated using the CRISPR/Cas9 system on C57BL/6J background, as described previously(Aizawa *et al*., under review). *Gsdmd*^–/–^*Gsdme*^–/–^ mice were developed by crossing *Gsdmd*^–/–^ mice with *Gsdme*^–/–^ mice. Only male mice were used to exclude the influences of the female hormonal cycle. Mice were housed (4/cage, RAIR HD ventilated Micro-Isolator Animal Housing Systems, Lab Products, Seaford, DE) in an environment maintained at 23 ± 2°C with *ad libitum* access to food and water under a 12-h light/dark cycle with lights on from 8:00 to 20:00. All animal experiments were approved by the Use and Care of Experimental Animals Committee of the Jichi Medical University Guide for Laboratory Animals and carried out in accordance with the Jichi Medical University guidelines (Permit No. 23053-02, 24010-03).

### Cell culture

For isolation of thioglycollate-elicited macrophages, mice were injected intraperitoneally with 2 mL of 4% Brewer thioglycollate medium (211716, Becton Dickinson, Franklin Lakes, NJ), and peritoneal cells were collected 3 days after injection. The isolated cells from each mouse were seeded at 1 x 10^6^ cells/mL in 10% fetal calf serum (FCS)/RPMI 1640 medium. After 3 h, non-adherent cells were washed out with phosphate-buffered saline (PBS), and adherent cells were used as peritoneal macrophages. LentiX293T (Takara Bio, Shiga, Japan) cells were cultured in Dulbecco’s modified Eagle’s medium (DMEM; Wako, Osaka, Japan) supplemented with 10% FCS, 1 mM sodium pyruvate, and antibiotics. Mouse immortalized bone-marrow-derived macrophages (iBMDMs) were gifted from Dr. Eicke Latz and cultured in DMEM supplemented with 10% FCS and antibiotics(Hornung *et al*, 2008). Unless otherwise indicated, cells were cultured at 37°C in 5% CO_2_.

### Plasmids

To develop DmrB-ASC or NINJ1-mNeonGreen-expressing lentiviral vector, PCR-generated DmrB, human ASC, human NINJ1, and mNeonGreen, were Gibson subcloned into CS-EF-1 or CS-IV-TRE-EF-1-KT (derived from CS-CA-MCS or CS-IV-TRE-CMV-KT; RIKEN BRC). The sgRNA targeting Ninj1, Xkr8, Ano6, and Tmem63b were designed with benchiling (https://www.benchling.com) and subcloned into LentiCRISPRv2, which was a gift from Feng Zhang (Addgene plasmid #52961; http://n2t.net/addgene: 52961; RRID: Addgene_52961). The gRNA targeting GSDMD and GSDME were developed using CRISPICK (https://portals.broadinstitute.org/gppx/crispick/public) and subcloned into pRDA052 expressing enCas12a(DeWeirdt *et al*, 2021). The sequence of the enCas12a gRNA targeting GSDMD and GSDME with DR2 is as follows: CATCGACGACATCAGAGACTTTGTAATTTCTACTATCGTAGATAAACGAAGAGTGACTCTCCACCA. The details of the sequences are described in the key resource table.

### Lentiviral preparation

LentiX293T cells were co-transfected with self-inactivating vectors, pLP1, pLP2, and pVSVG using PEI MAX (Polysciences, Warrington, PA) to prepare the lentiviral vectors. Culture media containing the lentiviral vectors were collected 3 days after transfection. The collected media were filtered with a 0.45-µm filter and ultracentrifuged at 21,000 rpm using a SW55 Ti rotor (Beckman Coulter, Brea, CA), and the pellets were resuspended in phosphate-buffered saline (PBS) containing 5% FCS. For lentiviral transduction, the cells were incubated with purified lentiviral vectors in the presence of 8 µg/mL polybrene (Sigma, St. Louis, MO). The details of the developed cells are described in the key resource table.

### Analysis of culture supernatants

Primary peritoneal macrophages isolated from WT, *Casp1/11*^–/–^, *Gsdmd*^–/–^, *Gsdme*^–/–^, and *Gsdmd*^–/–^*Gsdme*^–/–^ mice were seeded at 1.25×10^5^ cells/well into 96 well plates. After indicated treatments, culture supernatants were collected, and the IL-1α and IL-1β levels were measured by enzyme-linked immunosorbent assay (ELISA) using a commercial kit (R&D Systems, Minneapolis, MN, USA). LDH released from cells was measured by a Cytotoxicity Detection kit (Roche, Mannheim, Germany) according to the manufacturer’s instructions. To determine the total cellular LDH activity, cells were lysed with 2% TritonX100. L-Lactate Dehydrogenase from rabbit muscle (Roche) was used as a standard.

### Analysis of neutrophil infiltration

For administration of cholesterol crystal, crystals are resuspended in PBS at a concentration of 1 mg/mL and injected intraperitoneally (2 mg/mice). After 6 h, cells were collected from peritoneal lavage and analyzed using flow cytometry (BD FACS Lyric; BD Biosciences, San Jose, CA, USA). The cells were labeled with the following antibodies: fluorescein isothiocyanate (FITC)-conjugated anti-CD45R (11-0452, eBioscience, San Diego, CA), phycoerythrin (PE)-conjugated anti-Ly6G (561104, BD Biosciences), and allophycocyanin (APC)-conjugated anti-CD45 (17-0451, eBioscience). The data were processed by FlowJo software. Isotype control antibodies were used as negative controls to exclude nonspecific background staining.

### Live cell monitoring of SYTOX Green

Cells were incubated with 1 µg/mL Hoechst33342 (Dojindo, Kumamoto, Japan) for 20 min, and then cultured in the presence of 100 nM SYTOX Green (Thermo Fisher Scientific, Waltham, MA, USA) for 30 min. After labeling, cells were stimulated with the indicated reagents. Fluorescence intensity was measured by using a multimode microplate reader (Spark; TECAN, Männedorf, Switzerland).

### Live cell imaging

Peritoneal macrophages were seeded at 2.5 ×10^5^ cells on an 8-well coverglass chamber (IWAKI, Shizuoka, Japan) and primed with Pam3CSK4 for 18 h, then labeled with the indicated probes. To assess membrane permeability, cells were stained with Hoechst33342 for 20 min and maintained with medium containing 100 nM SYTOX Green. After images were obtained at 0 h, cells were stimulated with the indicated reagents and images were captured at the indicated time points using confocal microscopy (FLUOVIEW FV10i; Olympus, Tokyo, Japan). For analysis of cell morphology, primed cells were stained with 5 µM CFSE (Thermo Fisher) for 30 min in the presence of 0.04% Pluronic F127 (Sigma) and cultured in the medium containing 100 nM SYTOX Deep red. Z-stack time-lapse images were captured using confocal microscopy. The morphological parameters of the cells were analyzed using Imaris10.0 (Oxford Instruments, Abingdon-on-Thames, UK). For analysis of PtdSer, primed cells were stimulated with nigericin in DMEM without phenol red in the presence of FITC-Annexin V and DRAQ5 (Biolegend, San Diego, CA).

### Analysis of annexin V-positive cells

For flow cytometric analysis, iBMDMs were seeded on 12 well plates. After indicated treatments, cells were collected and stained with FITC-Annexin V and 7-AAD and analyzed by BD FACS Lyric. The data were processed by FlowJo software. For high content analysis, iBMDMs were seeded on Phenoplate 96-well plates (Revvity, Waltham, MA) and stained with Hoechst33342 for 20 min. Then, cells were stimulated with 100 nM B/B dimerizer in the presence of FITC-Annexin V and 100 nM SYTOX Deep red. After 2 h, cells were analyzed by an Operetta CLS high-content analysis system (PerkinElmer, Waltham, MA).

### Evaluation of membrane tension using Flipper-TR

Peritoneal macrophages were seeded at 2.5 ×10^5^ cells on an 8-well coverglass chamber and primed with Pam3CSK4 for overnight, then labeled with 1 µM Flipper-TR (Cytoskeleton, Denver, CO, USA) for 15 min. After labeling, cells were pretreated with 5 mM glycine for 30 min and stimulated with 5 µM nigericin for 30 min in the presence of SYTOX Deep red. Fluorescence lifetime imaging microscopy (FLIM) was performed using SP8 FALCON FLIM (Leica Microsystems).

### Western blot analysis

Samples were separated by sodium dodecyl sulfate-polyacrylamide electrophoresis (SDS-PAGE) and transferred to PVDF membranes. After blocked with Blocking One (NACALAI TESQUE, Kyoto, Japan) for 30 min, the membranes were incubated for overnight at 4°C with the following primary antibodies: anti-β actin, (A5441; Sigma), anti-caspase-1(AG-20B-0042-C100; Adipogen, Farmingdale, NY, USA), anti-caspase-3 (#9662; Cell Signaling Technology, Danvers, MA), anti-GSDMD (#50928; Cell Signaling Technology), anti-GSDMD (ab209845; Abcam, Cambridge, UK), anti-GSDME (ab215191; Abcam), anti-IL-1α (AF-400-SP; R&D Systems), anti-IL-1β (AF-401NA; R&D Systems), anti-NINJ1(clone 25; kindly gifted from Dr. Kayagaki, Genentech) (Kayagaki *et al*., 2021), and anti-NLRP3 (AG-20B-0014; Adipogen). As secondary antibodies, HRP-goat anti-mouse Superclonal IgG (Thermo Fisher Scientific), HRP-goat anti-rabbit IgG (Cell Signaling Technology), and HRP-rabbit anti-goat IgG (Thermo Fisher Scientific) were incubated with the membrane for 1 h. After washing with TBS-Tween, immunoreactive bands were visualized by Western Blot Quant HRP substrate (TAKARA Bio) or Western BLoT Ultra Sensitive HRP substrate (TAKARA Bio).

### Statistical analysis

Data are expressed as mean ± standard deviation (SD). Differences between the two groups were determined by Student’s t-test. Differences between multiple group means were determined by two-way analysis of variance (ANOVA) combined with Tukey’s post hoc test. Differences between multiple groups with repeated measurements were evaluated by repeated one-way ANOVA or repeated two-way ANOVA combined with the post hoc test. Analyses were performed using GraphPad Prism 10 software (Graph Pad Software, La Jolla, CA, USA). A p-value of < 0.05 was considered statistically significant. Biological replicates in primary culture indicate cells derived from each mouse. Biological replicates in cell line experiments indicate replicates of the same experiment conducted upon separately seeded cultures on separate days. The number of biological replicates is described in the figure legend.

## Data availability

The code used for data processing and image analysis (R scripts and FIJI macros) is available upon reasonable request.

## Author Contributions

Conceptualization, T.Ka., and M.T.; Methodology, T.Ka., Y.K., and T.Ku.; Validation, T.Ko. and Y.M.; Investigation, T.Ka., H.A., E.A., T.Ko., and C.B.; Resources, E.A., T.Ka; Writing- Original Draft, T.Ka.; Writing- Review & Editing, T.Ka., M.T.; Visualization, T.Ka.; Supervision, T.Ko. and T.M.; Project Administration, T.Ka., and M.T.; Funding Acquisition, T.Ka. and M.T.

## Acknowledgments

This study was supported by grants from the Japan Society for the Promotion of Science (JSPS) through Grants-in-Aid for Scientific Research (24K11219 to MT), the NAITO Foundation (T.Ka.) and INAMORI foundation (T.Ka.). We are grateful to Dr. Jun Suzuki (Kyoto University, Kyoto, Japan) and Dr. Tadashi Kasahara (Jichi Medical University) for his valuable suggestions. We appreciate Dr. Nobuhiko Kayagaki for providing essential materials and kind support (Genentech).

## Abbreviations

ASC: apoptosis-associated speck-like protein containing a caspase recruitment domain
DAMPs: damage/danger-associated molecular patterns
DKO: double-knockout
GSDMD: gasdermin D
GSDME: gasdermin E
iBMDM: immortalized bone marrow-derived macrophages
IL: interleukin
LDH: lactate dehydrogenase
NLRC4: nucleotide-binding oligomerization domain, leucine-rich repeat and caspase recruitment domain containing 4
NLRP3: nucleotide-binding oligomerization domain, leucine-rich repeat and pyrin domain
PRR: pattern recognition receptor
PtdSer: phosphatidylserine
RCD: regulated cell death
WT: wild-type

---

**Fig. S1.**
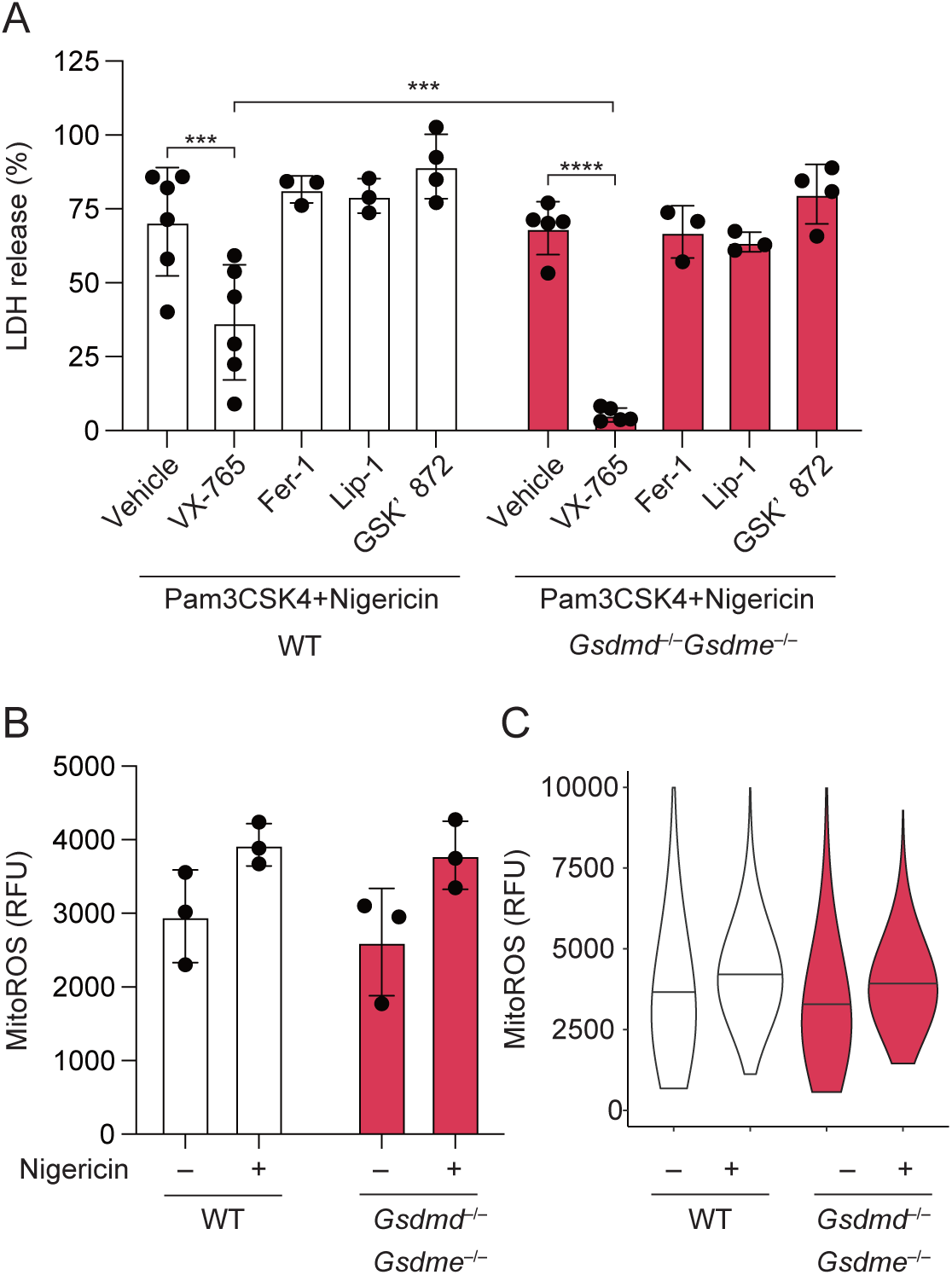
Necrotic cell death triggered by inflammasome is mediated by caspase-1. (A) Primed WT and *Gsdmd*^–/–^*Gsdme*^–/–^ macrophages were pretreated with indicated inhibitors and treated with nigericin for 3 h. The levels of LDH in the supernatants were assessed. (B and C) Primed WT and *Gsdmd*^–/–^*Gsdme*^–/–^ macrophages were labeled with MitoSOX and treated with nigericin for 2 h. (A and B) The data were obtained from cells derived from 3 – 6 mice per group. Each dot represents one mouse; the bar indicates mean ± SD. (C) Representative violin plot from cells of a single mouse.

**Fig. S2.**
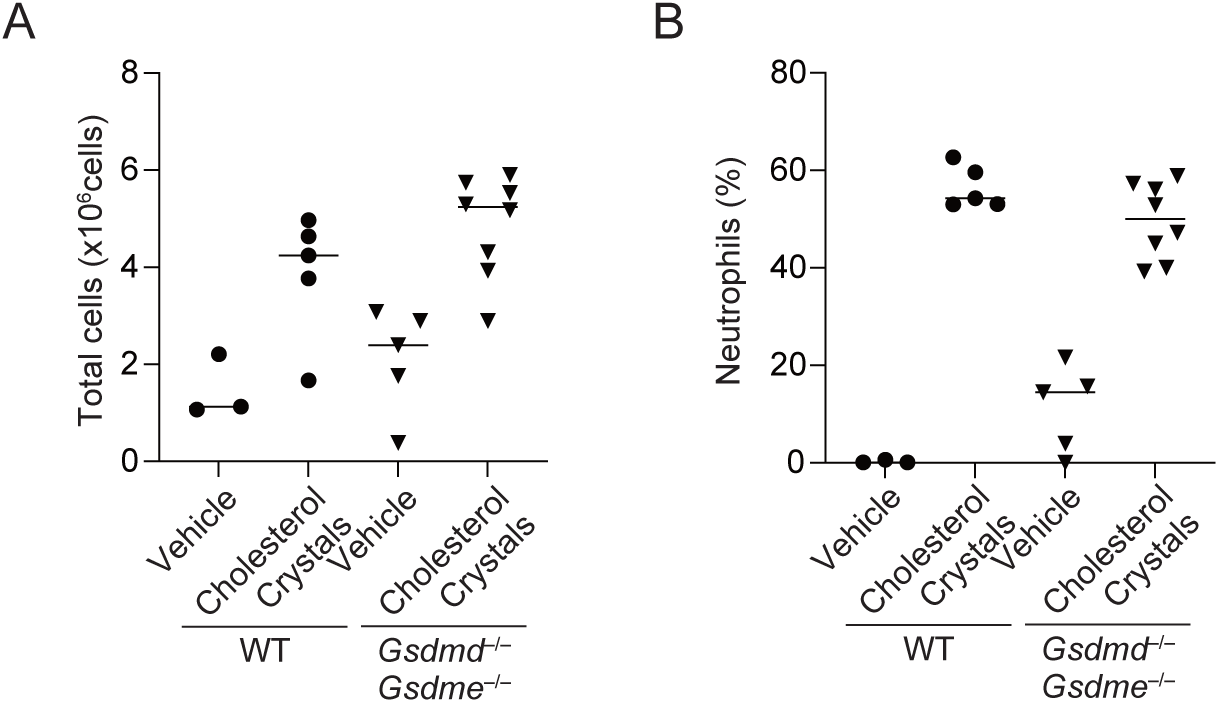
GSDMD and GSDME are dispensable for neutrophil infiltration induced by cholesterol crystals. (A and B) WT and *Gsdmd*^–/–^*Gsdme*^–/–^ mice were intraperitoneally administered cholesterol crystals. After 6 h, peritoneal cells were harvested and analyzed by flow cytometry. (A) Representative plots of total cells. (B) The ratio of infiltrated neutrophils. The data were obtained from 3–8 mice per group. Each dot represents one mouse; the bar indicates the median.

**Fig. S3.**
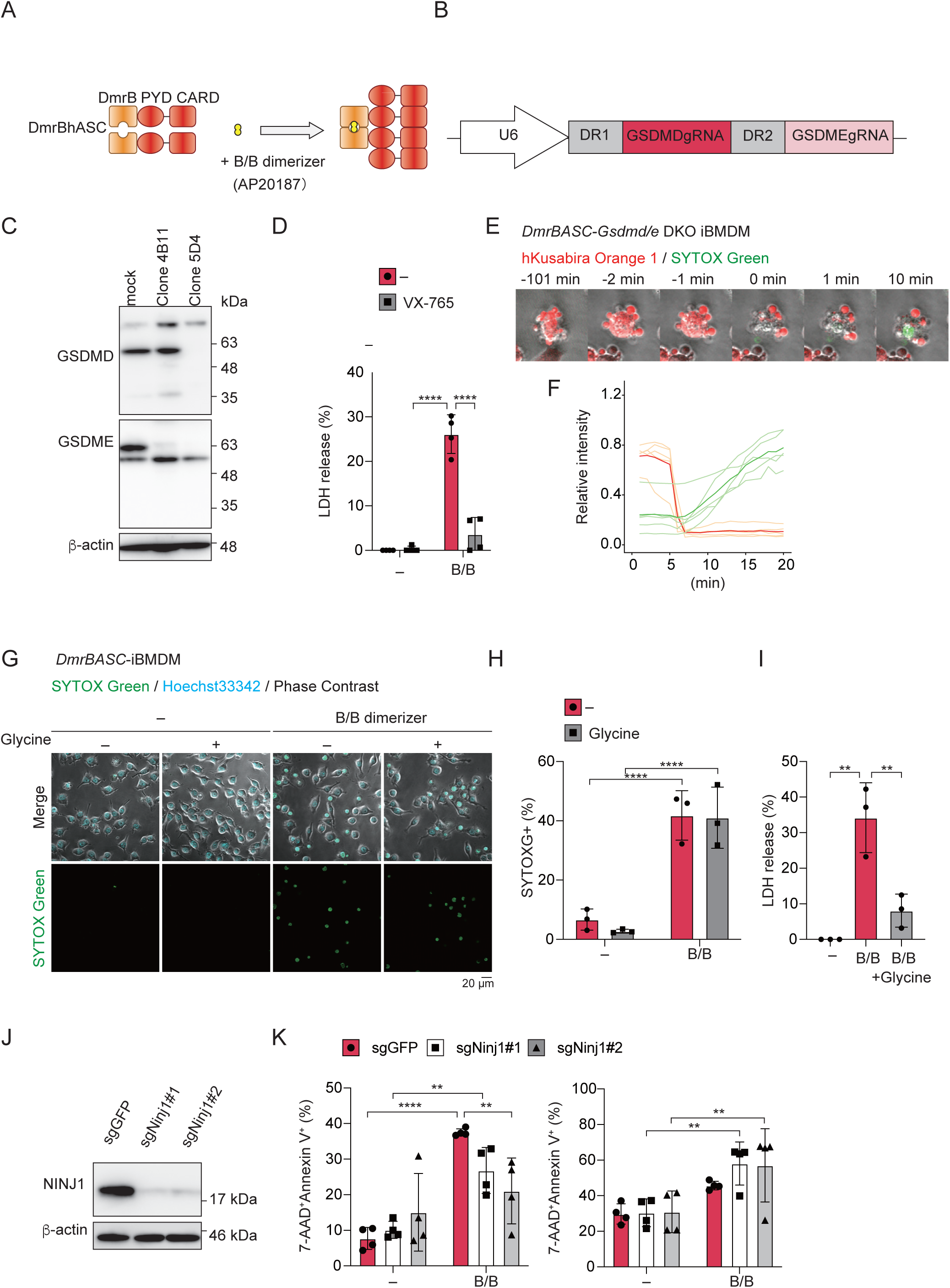
NINJ1-mediated plasma membrane rupture in DmrBASC–Gsdmd/e DKO iBMDM. (A) A schematic model of DmrBASC fusion protein. (B) The design of tandem gRNA targeting GSDMD and GSDME. (C) Protein from *DmrBASC*–*Gsdmd/e* DKO iBMDM were analyzed by western blot. (D) *DmrBASC*–*Gsdmd/e* DKO iBMDM were pretreated with VX-765 for 30 min and treated with 100 nM B/B dimerizer for 4 h. The levels of LDH in the supernatants were assessed. (E and F) *DmrBASC*–*Gsdmd/e* DKO iBMDM were transduced with lentiviral vector encoding Kusabira Orange and images were captured by confocal microscopy in the presence of SYTOX Green. (E) Montage images and (F) quantification of fluorescent signals. (G–I) *DmrBASC*–iBMDM were pre-treated with 5 mM glycine and treated with 100 nM B/B dimerizer in the presence (G and H) or absence of SYTOX Green (I). (G and H) Images were visualized by confocal microscopy. (H) The percentage of SYTOX Green-positive cells was quantified. (I) The levels of LDH were assessed at 4 h. (J and K) *DmrBASC*–*Gsdmd/e* DKO iBMDM were transduced with LentiCRISPRv2 expressing GFP or *Ninj1*-targeted gRNA. (J) NINJ1 protein expression was analyzed by western blot. (K) Cells were treated with100 nM B/B dimerizer for 4 h and stained with FITC-Annexin V and 7-AAD for flowcytometric analysis. (D, G, H, I, and K) The data were obtained from 3 independent experiments. Each dot represents independent experiments; bar indicates mean ± SD. (E and F) The data were obtained from 2 independent timelapse imaging. Light orange lines and light green lines indicate individual cells, and the deep orange line and deep green line indicates the mean of cells. Statistical significance was calculated using two-way ANOVA with Tukey’s post hoc test. \*\**p* < 0.01, \*\*\*\**p* < 0.0001.

**Fig. S4.**
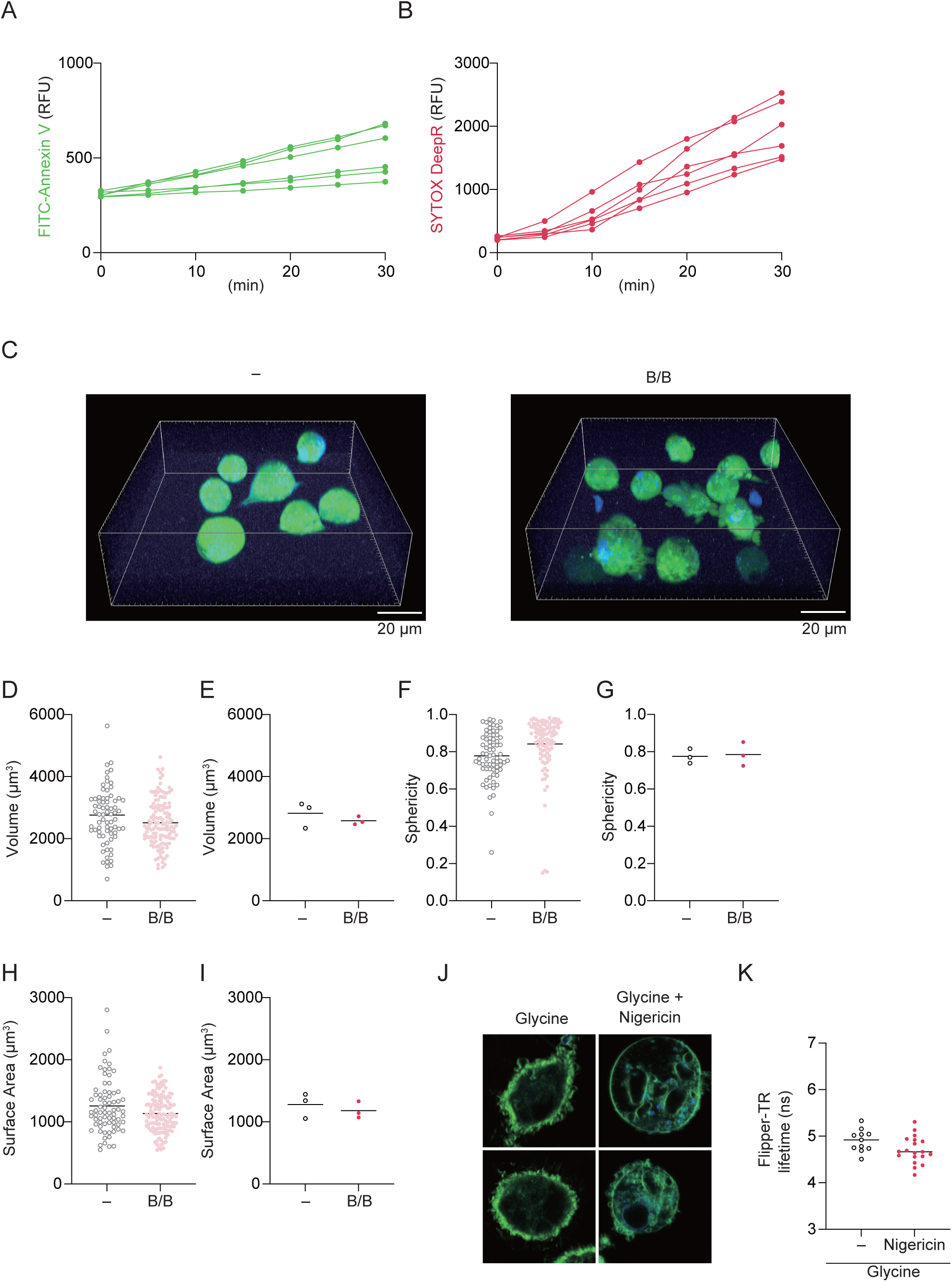
Exposure of phosphatidyl serine during inflammasome-driven necrotic cell death. (A and B) Primary peritoneal macrophages isolated from *Gsdmd*^–/–^*Gsdme*^–/–^ mice were primed with Pam3CSK4 (100 ng/mL) for overnight and treated with 5 µM nigericin in the presence of FITC-Annexin V and SYTOX Deep red. Z-stack timelapse imaging was performed at 5 min interval. Annexin V-positive cells were tracked and analyzed. T_0_ is defined as a time frame in which FITC-Annexin V signals was detected. Quantification of fluorescent signals of Annexin V (A) and SYTOX Deep red (B). Each dot represents medians of all cells obtained from single mouse (N=6). (C–I) *DmrBASC*–*Gsdmd/e* DKO iBMDM were stained with 5 µM CFSE and treated with100 nM B/B dimerizer for 30 min. Z-stack imaging was performed. (C and D) Representative 3D images of unstimulated cells (C) and cells (D) treated with B/B dimerizer. The cell volume (D and E), sphericity (F and G), and surface area (H and I) were analyzed. (D, F, and H) Each dot represents an individual cell from three independent experiments. (E, G, I) Each dot represents mean value per experiment. (J and K) Primary peritoneal macrophages isolated from *Gsdmd*^–/–^*Gsdme*^–/–^ mice were primed with Pam3CSK4 (100 ng/mL) for overnight and labeled with Flipper-TR for 15 min. The labeled cells were pretreated with 5 mM glycine and stimulated with 5 µM nigericin for 30 min. Fluorescent lifetime was measured using FLIM. (J) Representative FLIM imaging and (K) lifetime of Flipper-TR. Each dot represents individual cell.

**Fig. S5.**
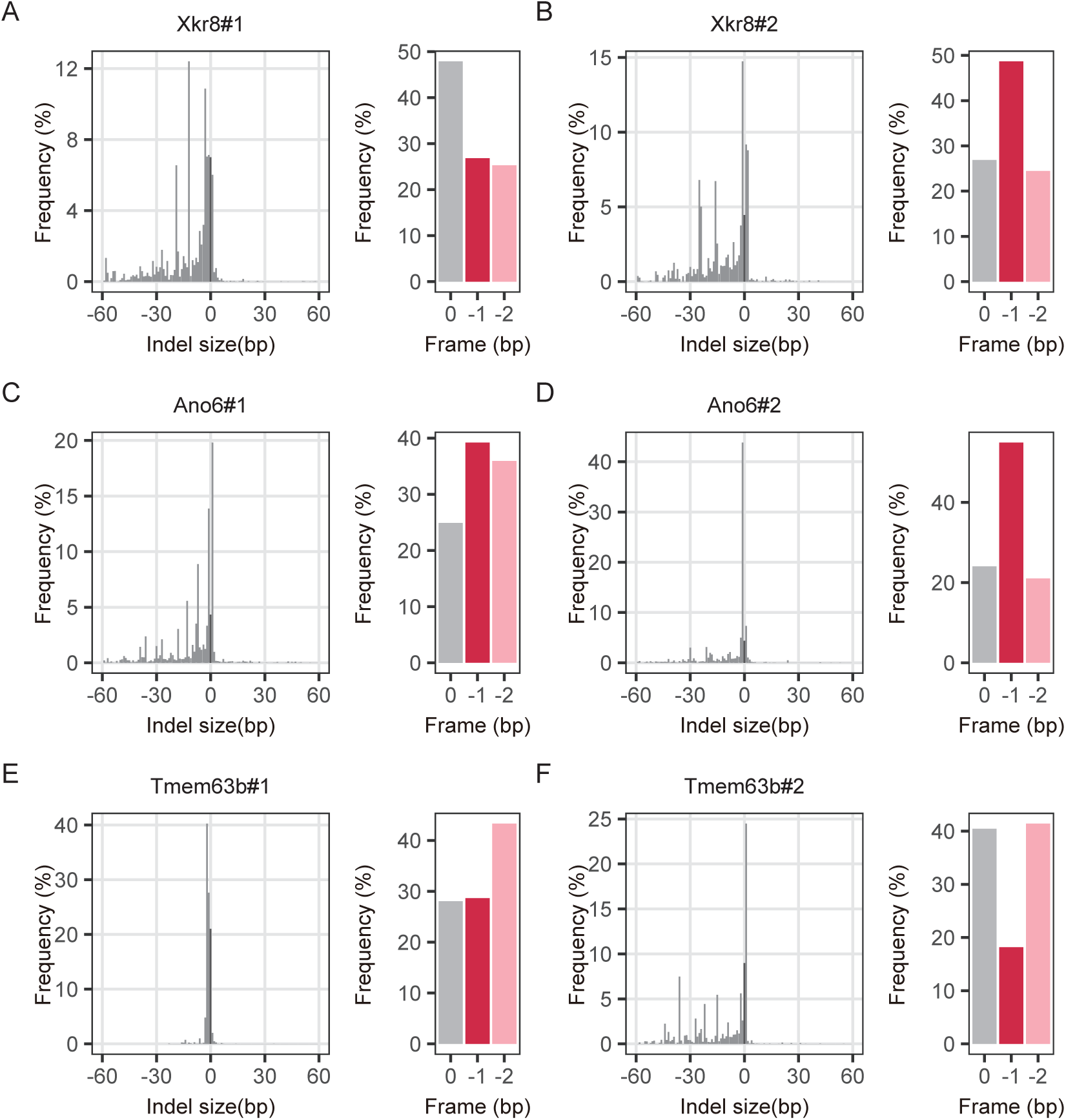
Development of DmrBASC–Gsdmd/e DKO iBMDM lacking phospholipid scramblases. (A–F) *DmrBASC*–*Gsdmd/e* DKO iBMDM were transduced with LentiCRISPRv2 expressing *GFP*, *Xkr8*, *Ano6*, and *Tmem63b*-targeted gRNA. The frequency of indel in *Xkr8*-(A and B), *Ano6*-(C and D), and *Tmem63b* (E and F)*-*ablated iBMDM was analyzed by amplicon sequencing and subsequent CRISPResso2 analysis.

**Fig. S6.**
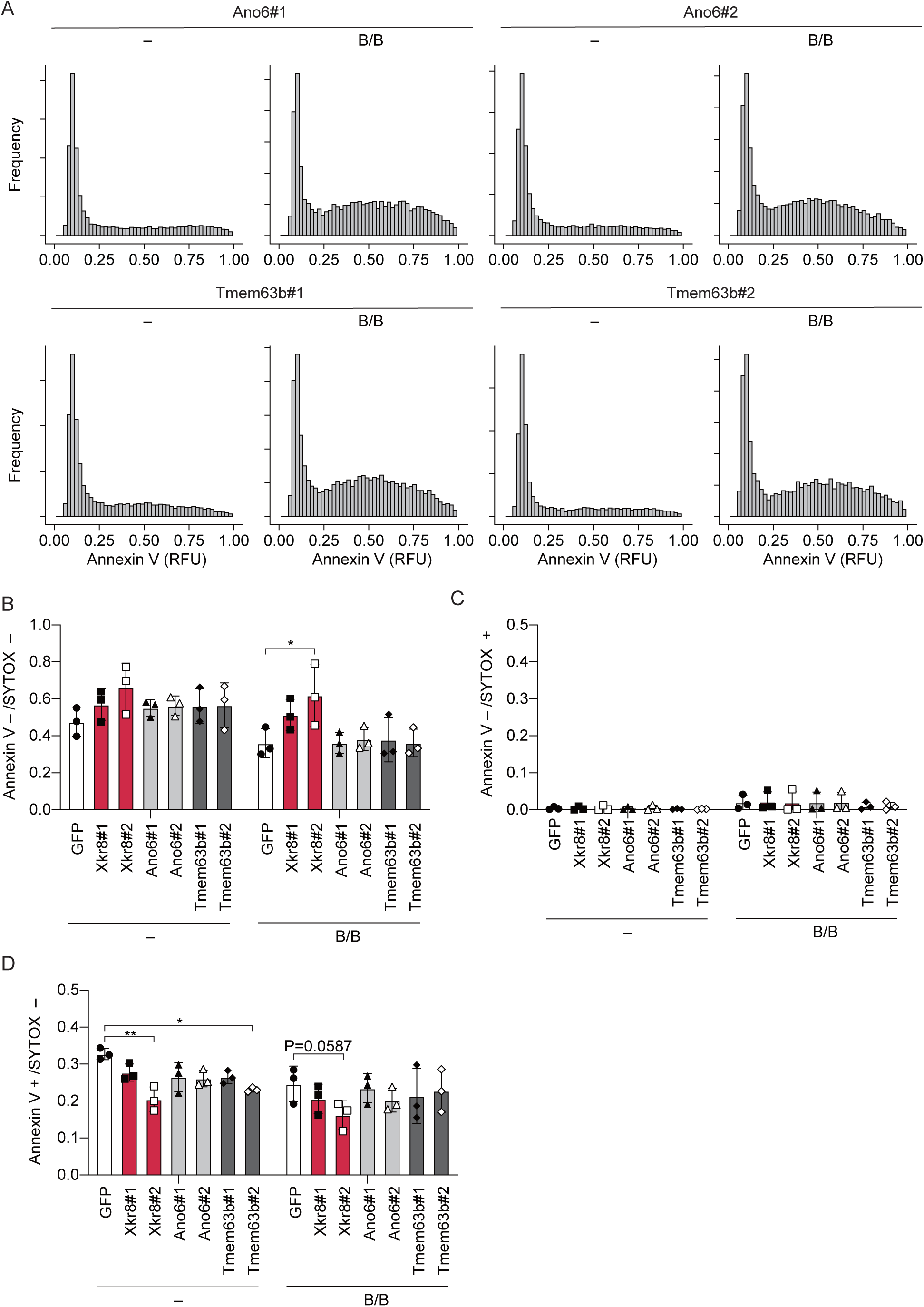
PtdSer exposure in DmrBASC–Gsdmd/e DKO iBMDM lacking phospholipid scramblases. (A–F) *DmrBASC*–*Gsdmd/e* DKO iBMDM were transduced with LentiCRISPRv2 expressing *GFP*, *Xkr8*, *Ano6*, and *Tmem63b*-targeted gRNA. The cells were treated with 100 nM B/B dimerizer. (A) Representative histogram of Annexin V intensity in cells expressing gRNA targeting *Ano6* and *Tmem63b*. (B–D) The ratio of Annexin V– SYTOX deep red– cells (B), Annexin V– SYTOX deep red+ cells (C), and Annexin V+ SYTOX deep red– cells (D) were quantified. (A–D) The data are obtained from 3 independent experiments. (B–D) Each dot represents individual experiments; the bar indicates mean ± SD. Statistical significance was calculated using two-way ANOVA with Tukey’s post hoc test.

## Notes

### Competing Interest Statement

The authors have declared no competing interest.

